# Improved detection and spatiotemporal spectral analysis of neural traveling waves

**DOI:** 10.64898/2026.09.11.750831

**Authors:** Martin Vinck, Carmen Gasco-Galvez, Miguel Rodrigues, Jakob C. B. Schwenk, Misako Komatsu, Frederic Chavane, Andres Canales-Johnson, Andrea Alamia

## Abstract

Traveling waves (TWs) are a fundamental mode of neural dynamics, yet existing detection methods are limited by sensor geometry, spatial-frequency resolution, signal amplitude, and ambiguity between propagating and standing-wave patterns. Here we introduce the Traveling Wave Index (TWINDEX), a framework for three-dimensional spatiotemporal spectral analysis of TWs across temporal frequency, spatial frequency, and propagation direction. TWINDEX generalizes to irregular sensor layouts, and quantifies wave strength as the reduction in circular phase variance produced by a candidate planar wave. This normalization yields robust behavior at both low and high spatial frequencies and suppresses coherent in-phase activity. Directional moments further separate planar from standing waves. We derive analytical links between TWINDEX, parametric planar-wave fit, and distance-phase correlation, and introduce projected distance-phase correlation (ProDPC) for sensitive single-trial planar-wave detection. Applying these methods to large-scale marmoset ECoG and human EEG, we identify alpha/low-beta TWs localized in spatial and temporal frequency, with physiologically plausible propagation speeds. Across trials and epochs, waves occur in oppositely directed propagation modes, demonstrating that alpha/beta activity is associated with both feedforward- and feedback-directed large-scale dynamics.

## 1. Introduction

The past decades have seen numerous discoveries of distinct traveling waves (TW) patterns in large-scale recordings of cortical activity, with systematic associations to cognitive functions and brain state (Rubino et al., 2006; Zanos et al., 2015; Benucci et al., 2007; Nauhaus et al., 2012; Das et al., 2022; Zhang et al., 2018; Alexander et al., 2019; Nunez and Srinivasan, 2006; Chemla et al., 2019). Traveling waves are increasingly recognized as a fundamental organizing principle of brain activity, coordinating neural processing across space and time (Cruddas et al., 2025; Muller et al., 2018; Sato et al., 2012; Ermentrout and Kleinfeld, 2001; Dugué and Chavane, 2025), and linking micro- to meso- to macro-scales.

TW detection methods can generally be divided into two kinds of methods. Single-trial detection methods fit specific relations between distance and phase (Muller et al., 2014), or fit a certain traveling wave pattern with some model’s parameters (Zhang et al., 2018; Townsend and Gong, 2018; Denker et al., 2018). These methods are typically used in single trials and have been predominantly applied to data with a high signal-to-noise ratio (i.e., intracortical recordings). However, a disadvantage of these methods is that they require a specific template and a single set of parameters (e.g., spatial or temporal frequency, wave direction, wave origin). A different approach relies on decomposition methods that combine data across trials and characterize the entire spectrum by decomposing the signal into a series of waves, or principal components (Alexander et al., 2016, 2019; Alamia and VanRullen, 2019; Ambrogioni et al., 2017). Such decomposition can address crucial questions, e.g. whether traveling waves occur at specific spatial frequencies, or rather have energy in a broad range of spatial frequency, resulting from e.g. diffusive flow. For example, in scalp-EEG recordings in humans, the 2-D FFT has been used to characterize the full spectral activity within a given time window and to distinguish between forward and backward waves across different frequency bands (Alamia and VanRullen, 2019; Luo and Ester, 2025). However, this method has several limitations: it operates in only one spatial dimension, does not apply to irregular/non-rectangular arrays, does not yield an interpretable measure of wave strength, and has a resolution limited by the Discrete Fourier Transform.

Here, we address these limitations by introducing a spectral decomposition framework for traveling waves, the generalized *k*–*f* spectrum. This framework generalizes beyond previous two-dimensional formulations and is applicable to irregular electrode grids. Based on this spectrum, we define a novel measure called the Traveling Wave Index (TWINDEX), a dimensionless measure of traveling wave strength with several key advantages. We apply the TWINDEX to previously published marmoset ECoG recordings (Canales-Johnson et al., 2021; Gelens et al., 2024) and human scalp-EEG (Pang et al., 2020), enabling a detailed characterization of how rhythmic brain activity propagates across cortical space.

## Results

### The generalized k–f spectrum

We first define a generalized *k*–*f* spectrum (gKFS), a representation of TW activity as a function of temporal frequency *f*, angular spatial frequency *k*, and wave direction *θ* (see Methods for details, for an overview of all methods see Table 1). The gKFS is designed to apply to arbitrary sensor geometries, including irregularly sampled and non-rectangular arrays.

**Table 1:**
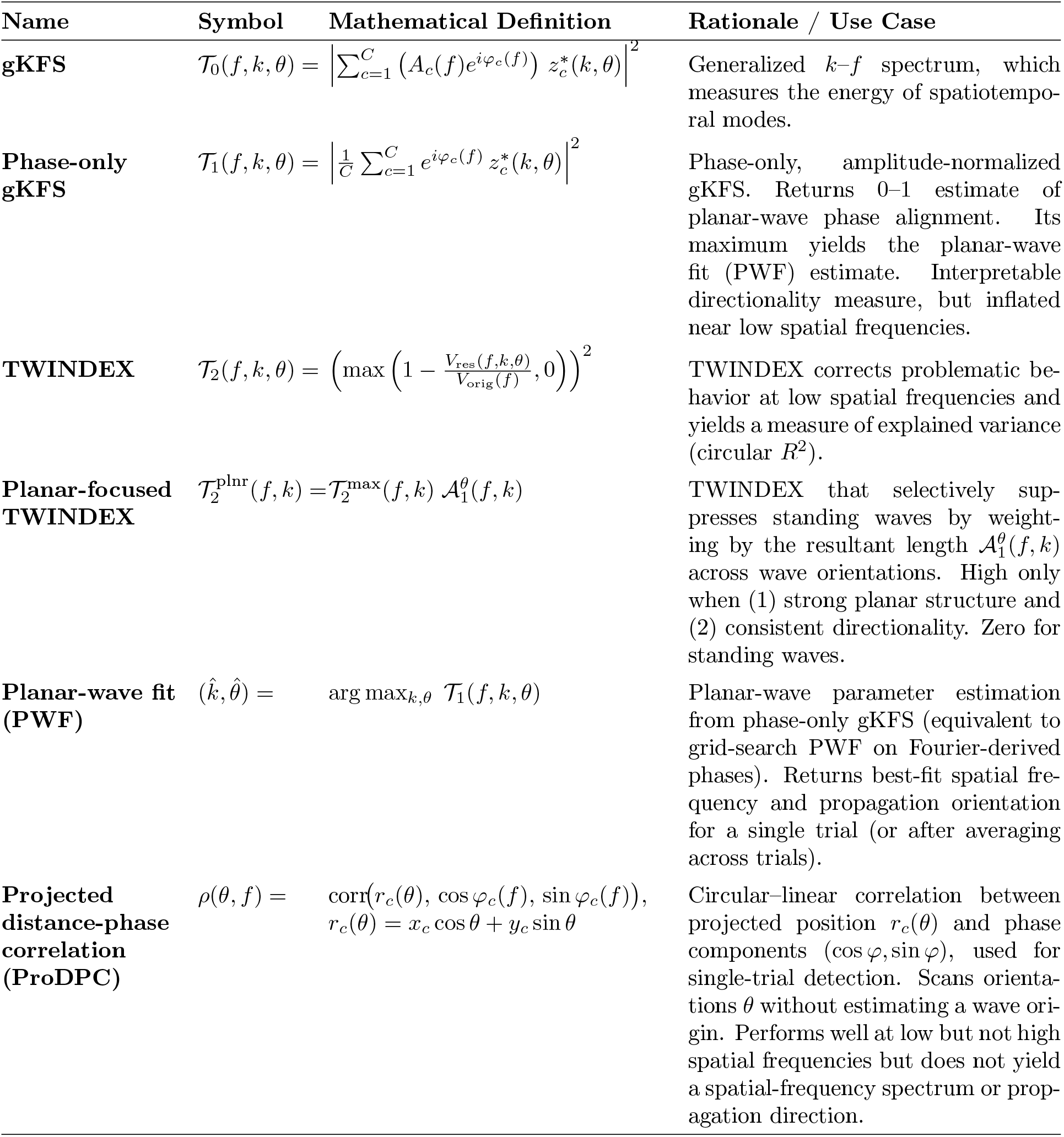
Overview of generalized *k*–*f* spectrum (gKFS)–based measures, their symbols, definitions, and conceptual motivations. All measures are computed per trial; dependence on trial index *p* is omitted.

To compute the gKFS, we first represent the signal at each channel in the frequency domain, for example using a Fourier transform, wavelet transform, or band-pass filtering followed by the Hilbert transform (Figure 1). For each temporal frequency *f*, spatial frequency *k* and wave orientation *θ*, we then evaluate the match between the spatial map of complex-valued signal representation and the planar-wave templates (Figure 1A-B) (see Methods). This match is computed as the inner product of the complex phasors, times the amplitude spectrum (see Methods). Scanning across (*k, θ*) yields the gKFS, T_0_(*f, k, θ*) (Eq. 14), which quantifies the strength of the match between the observed spatial signal pattern and each candidate planar wave (Figure 1A-B).

**Figure 1:**
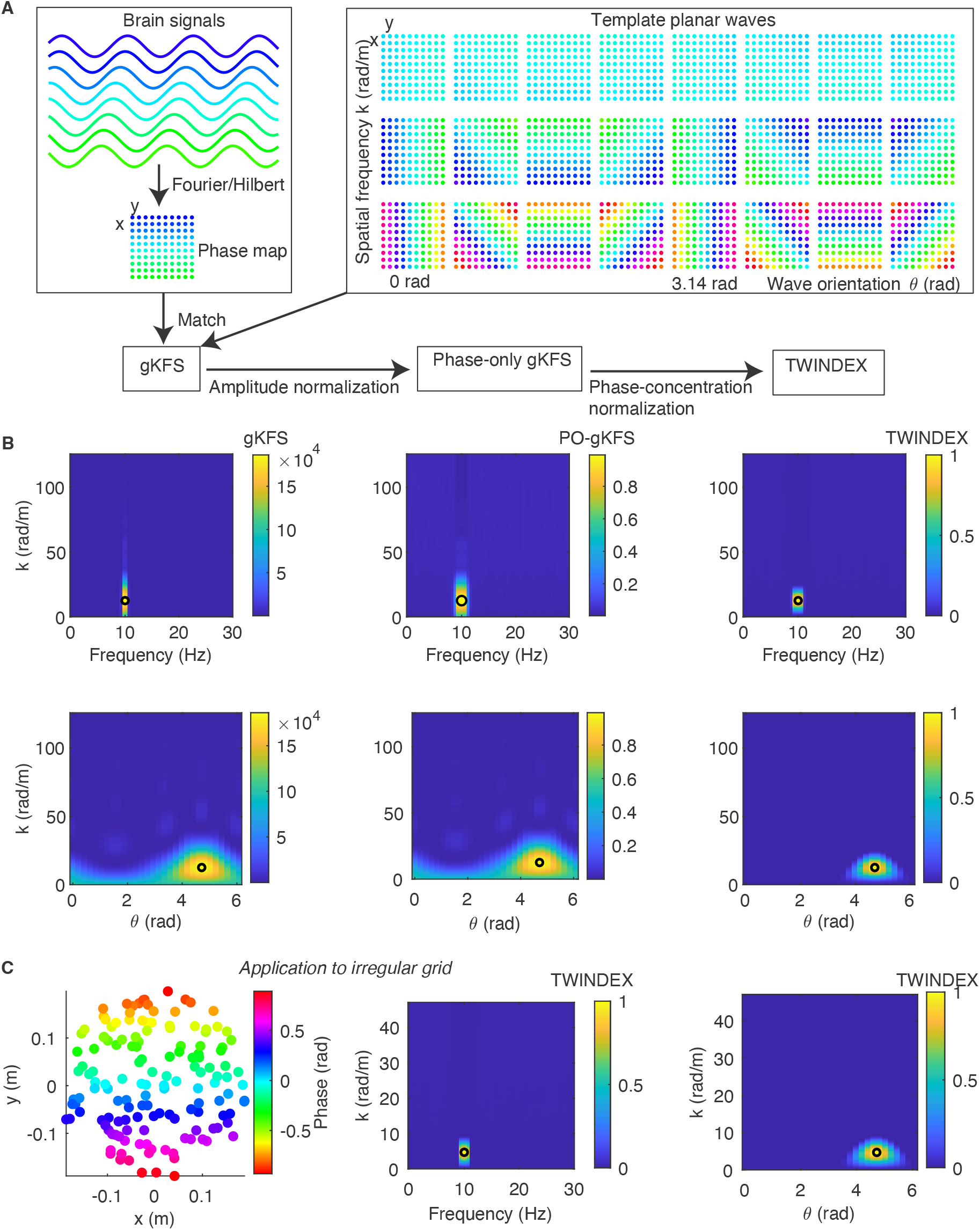
Illustration of the TWINDEX method. (A) Illustration of the steps to compute the gKFS (generalized *k*–*f* spectrum), phase-only gKFS, and the TWINDEX (see Results text / Methods). In brief, TWINDEX can be computed by scanning through planar waves of different spatial frequencies *k* and wave orientations *θ*, computing the match (inner product) with the signal Fourier Transform (yielding the gKFS), normalizing for amplitude (phase-only gKFS), and normalizing for the circular variance of the phase map to avoid artificial inflation towards low spatial frequencies (yielding the TWINDEX). (B) Top, Left-to-right: gKFS, phase-only gKFS, and TWINDEX as a function of temporal and spatial frequency. Black circle shows true parameters. Bottom: wave direction vs. spatial frequency. (C) Application of TWINDEX to irregular grids. Note that in this case, the template waveforms would be similar to (A) but be defined for an irregular grid.

The gKFS is related to a spatial Discrete Fourier Transform (DFT), but differs in two important aspects. First, the planar-wave kernels can be constructed for irregularly sampled electrode grids. Second, *k* is not restricted to the discrete Fourier frequencies of the recording aperture. For an array of spatial extent *L*, the spacing between DFT wavenumbers is 2*π/L* rad/m, corresponding to one complete spatial cycle across *L*. The gKFS can instead be evaluated on a densely interval of spatial frequencies, including *k <* 2*π/L* radians/m. This allows to detect TWs with low spatial frequencies, although the precision with which nearby spatial frequencies can be distinguished remains limited by the spatial aperture.

In simulations, the gKFS exhibits a peak at the simulated spatial frequency, temporal frequency, and wave orientation, including when the spatial phase gradient spans less than one cycle across the recording array (Figure S1D). The gKFS also recovers the correct wave parameters for irregular sensor layouts (e.g. irregularly sampled, not rectangular or square), for which a regular DFT cannot be used (Figure 1C).

### The phase-only generalized k–f spectrum

The gKFS is amplitude-sensitive, such that channels and frequencies with larger spectral amplitudes contribute more strongly to the spectrum. Consequently, the magnitude of the gKFS cannot be interpreted as a normalized measure of traveling-wave strength. For example, even when phases are randomly distributed across channels, the gKFS can exhibit large values at temporal frequencies with strong oscillatory power (Figure S2).

To remove this amplitude dependence, we normalize the complex-valued spectrum for each channel to unit magnitude, keeping only its phase (see Methods). We then evaluate the spatial match between this phase-only signal map and candidate planar waves with temporal frequency *f*, spatial frequency *k*, and wave orientation *θ* (Figure 1). This yields the phase-only generalized *k*–*f* spectrum (POgKFS), *T*_1_(*f, k, θ*) (Eq. 19). The PO-gKFS is bounded between 0 and 1 and quantifies the match between the phase map and candidate planar waves.

### The TWINDEX

Crucially, the ability of the PO-gKFS to characterize TWs deteriorates when the spatial frequency is low, resulting in a small phase gradient across the sensor array. For a planar wave with a spatial frequency *k*, the phase range across an array of length *L* is approximately *kL* radians. Thus, when *kL* ≪ 2*π* radians, the phases vary only weakly across sensors. We refer to this problem as the **TW aperture problem**. Low spatial frequencies can be expected to occur under various conditions in neural recordings (see Discussion): For a wave with temporal frequency *f* and propagation velocity *v*, the angular spatial frequency is

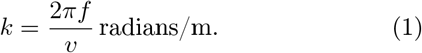

For example, a 10 Hz wave propagating at 1 m/s has a spatial frequency of approximately 10 cycles/m (62.8 rad/m), corresponding to a wavelength of 0.1 m. Across a recording array with a spatial extent of for example 0.02 m, as is typical for large-scale cortical recordings in smaller animals like marmosets or smaller grids in e.g. macaques, the resulting phase difference is only about 0.2 cycles. By contrast, for a larger recording array in humans of 0.3 m, the resulting phase difference constitutes 3 cycles, which would be a high spatial frequency.

We find that in the low-*k* regime, the PO-gKFS becomes poorly discriminative. The reason is that the POgKFS computes a match between the signal phase map and the candidate planar wave, effectively as a resultant vector length. This resultant vector length will be large in a wide range of low spatial frequencies, and at all wave orientations, if the phase map contains a narrow range of phases (see Methods). Consequently, the PO-gKFS will show high values near 1 for a broad range of spatial frequencies (including *k* = 0 and across all wave orientations (see Methods). We illustrate the TW aperture problem in simulations in Figure 2.

**Figure 2:**
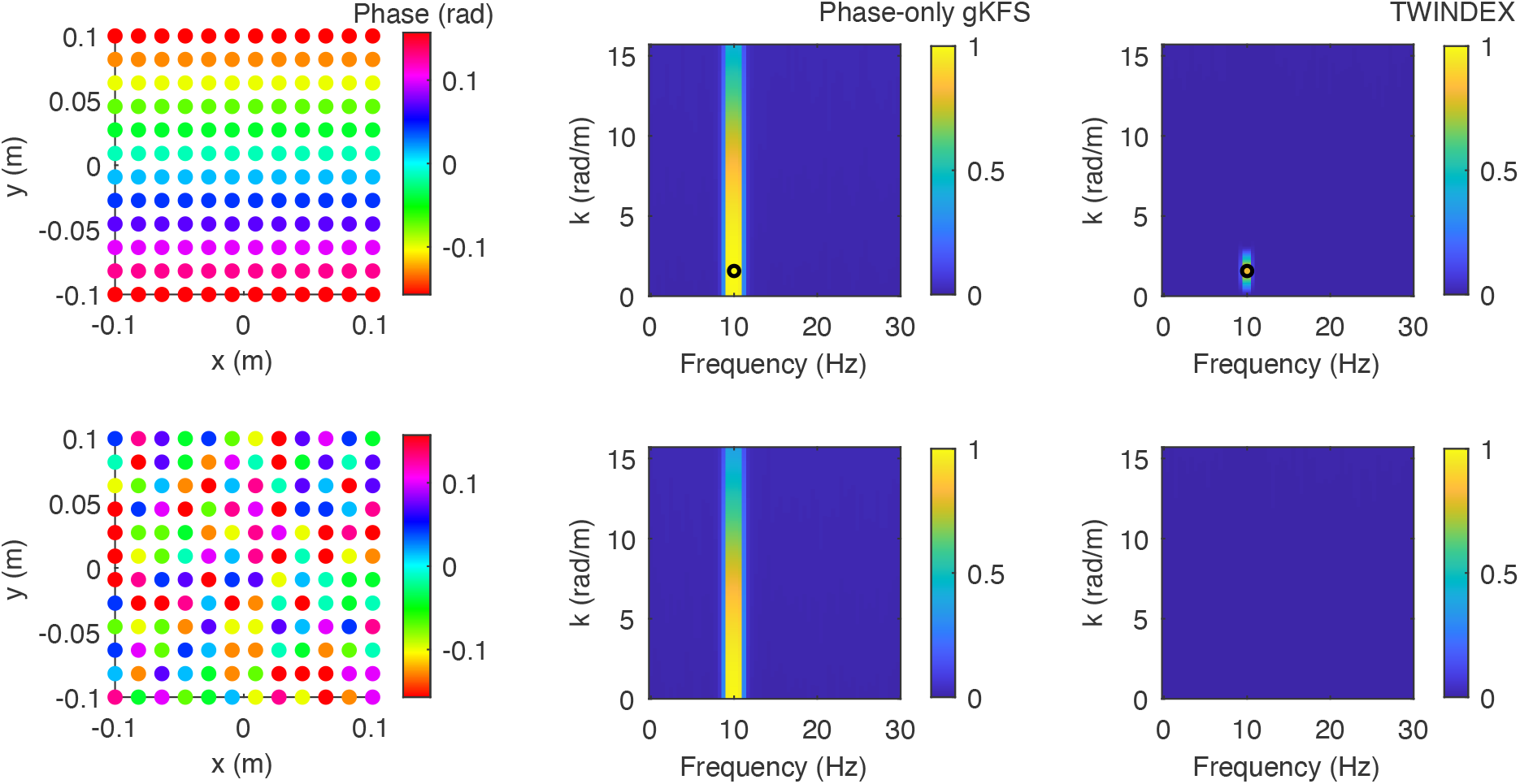
Comparison of Phase-only gKFS (PO-gKFS) and the TWINDEX. We show the case of planar wave (top row) or a random distribution of phases (bottom row). In both cases, an aperture problem occurs, as the spatial frequency is low and the range of phases is much smaller than 2 *π* radians. The TWINDEX correctly distinguishes the case of a genuine wave at low spatial frequency vs. a noisy organization of channel phases. By contrast, the PO-gKFS does not distinguish these two cases and has inflated values near *k* = 0. Black circle indicates true wave parameters.

To address the TW aperture problem, we introduce the Traveling Wave Index (TWINDEX), *T*_2_(*f, k, θ*) (Eq. 32; see Methods; Figure 1B), which can also be readily applied to irregular grids (Figure 1C). Like the PO-gKFS, the TWINDEX quantifies the match between the phase map and candidate planar waves. In addition to the POgKFS, the TWINDEX normalizes by the original circular variance (i.e. phase concentration) in the signal phase map. This normalization corrects for phase concentration that is already present in the original data and removes the uninformative *k* = 0 component. Importantly, this does not lead to a susceptibility to noise (see Figure 8).

To compare the TWINDEX and PO-gKFS, we simulated phase maps generated by low-spatial-frequency planar waves, but also by randomly shuffling the phases across channels. In simulations of low-spatial-frequency planar waves, TWINDEX exhibits a localized peak at the ground-truth spatial frequency, whereas the PO-gKFS shows a substantially broader and less discriminative spectrum (Figure 2). For randomly shuffled phases, we observed high PO-gKFS values across a broad range of *k*, whereas the TWINDEX showed values close to zero.

Importantly, as *k* → 0, the PO-gKFS will converge to a value of 1, while the TWINDEX fully suppresses a *k* = 0 component (see Methods for an analytical proof). This is highly relevant for electrophysiological recordings, where volume conduction (which is instantaneous in neural tissue of a common neural or noise source) can contribute *k* = 0 components (Vinck et al., 2011; Pesaran et al., 2018) (see further below for an empirical demonstration).

### The planar-focused TWINDEX

In neural recordings, spatial phase relationships may reflect not only planar TWs but also standing-wave patterns (Benucci et al., 2007). Standing wave patterns can arise from the spatial mixing of electric current dipoles through volume conduction (Pesaran et al., 2018) or the superposition of two counter-propagating planar waves. Standing-wave patterns can be identified from the angular structure of the TWINDEX spectrum (Figure 3): An ideal standing wave produces two opposing peaks separated by *π* radians.

**Figure 3:**
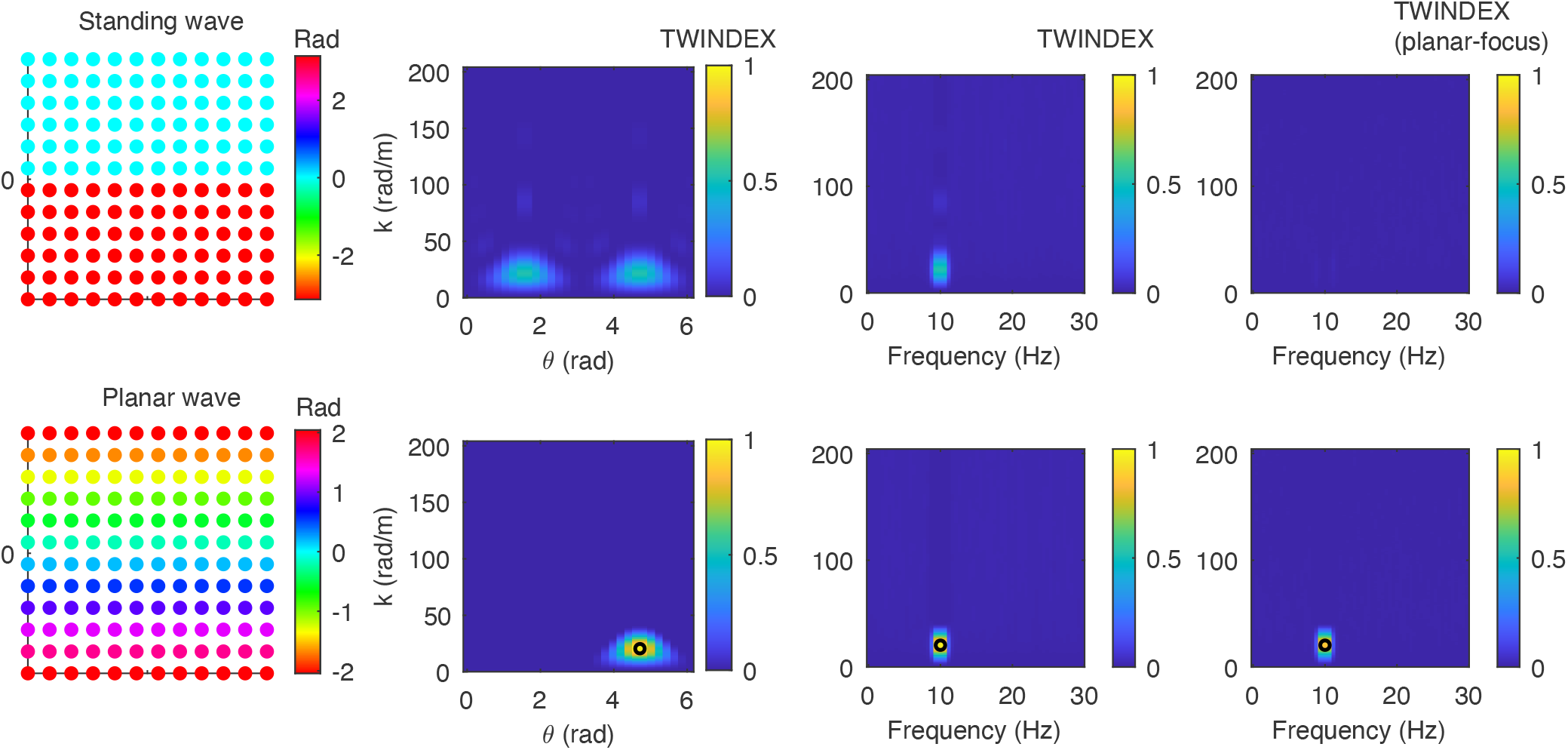
Planar-focused TWINDEX to dissociate standing and planar waves. Top row: case of a standing wave, which results in a bimodal TWINDEX for opposite wave orientations, and a corresponding peak at intermediate spatial frequencies in the TWINDEX. The planar-focused TWINDEX prevents the standing wave from appearing as a peak in the spectrum. Bottom row: planar wave, which results in a unimodal TWINDEX. In this case, the planar-focused TWINDEX correctly shows a peak. Thus, the planar-focused TWINDEX selectively picks up planar waves.

We quantify this directional asymmetry using the first circular moment *A*_1_ of the angular TWINDEX distribution (Eq. 42; see Methods). A large *A*_1_ indicates a preferential propagation direction, as expected for a planar TW, whereas a small *A*_1_ indicates a more directionally symmetric pattern. To distinguish standing waves from more isotropic angular patterns, we additionally use the second circular moment *A*_2_ (Eq. 43), which measures axial structure. Thus, a standing-wave pattern is characterized by low *A*_1_ but high *A*_2_, whereas a planar TW typically exhibits high values of both moments. Combining these angular moments with the magnitude of the TWINDEX yields wave-geometry-focused measures (Eqs. 44, 45; see Methods).

As illustrated in Figure 3, the planar-focused TWINDEX exhibits a clear peak for a simulated planar TW while being strongly suppressed for a standing wave.

### Traveling waves in large-scale marmoset data

Having established the theoretical properties of the TWINDEX, we demonstrate its utility by applying it to real neural recordings, illustrating how the method can characterize TWs in experimental data. We further analyzed data from two marmosets, in which ECoG recordings were made from the entire hemisphere. Marmosets engaged in a passive auditory listening task. We analyzed the pre-stimulus period.

We found that the data contained oscillations at 12.5 Hz (typical alpha / low-beta range) frequencies, with clear narrow-band spectral peaks in the power spectra (Figure 4A). In both monkeys, these 12.5 Hz waves were most prominent around the parietal cortex, and weakest in the occipital lobe (Figure 4B).

**Figure 4:**
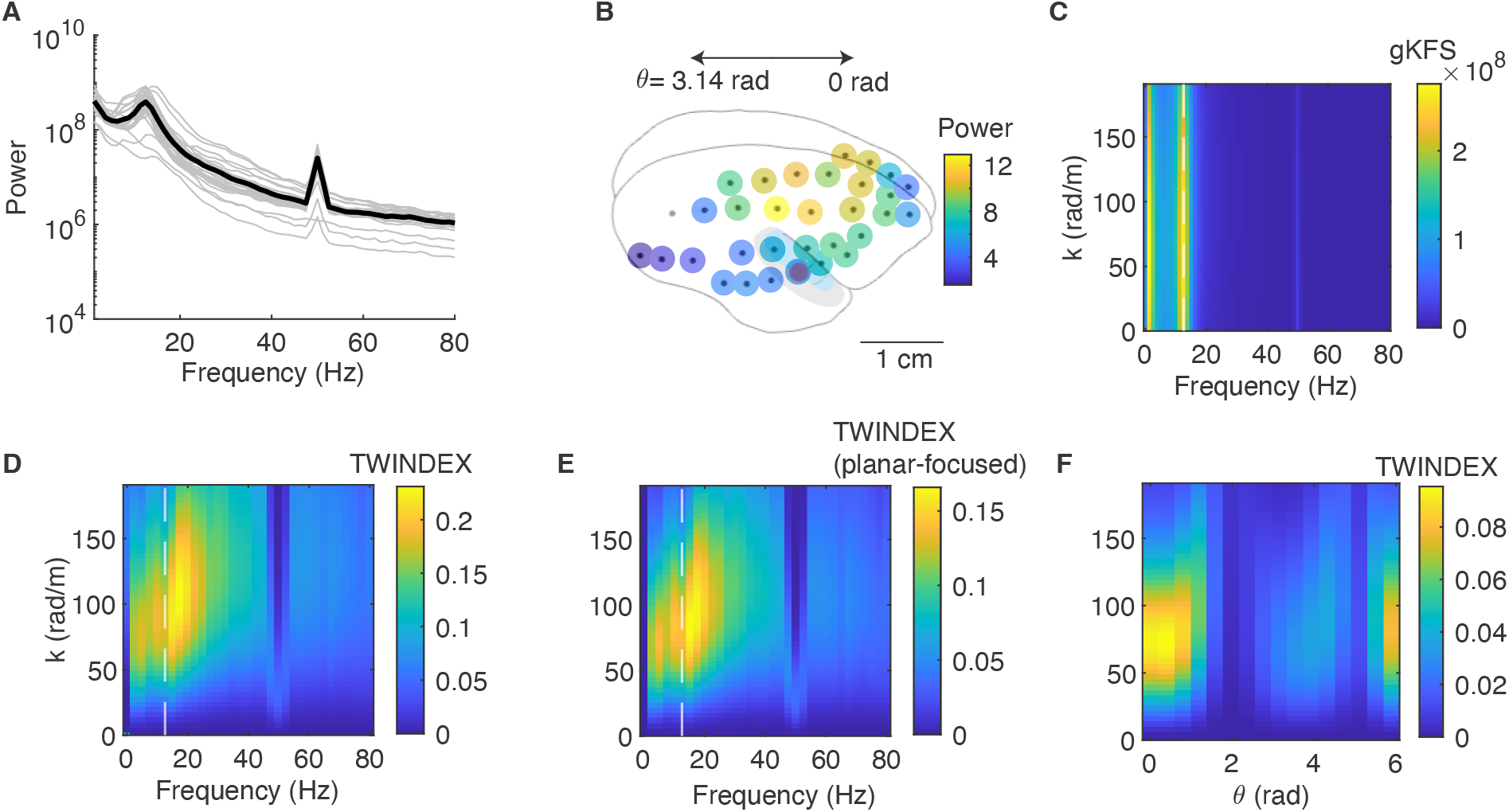
Analysis of ECoG marmoset data during pre-stimulus period, for monkey FR. (A) Power spectra across all channels. Note a genuine oscillatory band around 12.5 Hz, and line noise around 50 Hz. (B) Distribution of power (log-scaled) across channels. Note the shading of regions around the temporal area, containing auditory areas. (C) Average gKFS spectrum. (D) Average TWINDEX. White line indicates peak in power spectrum at 12.5 Hz. (E) Average planar-focused TWINDEX. (F) TWINDEX as a function of wave orientation and spatial frequency, at 12.5 Hz

We computed the gKFS, PO-gKFS and the TWINDEX (Figure 4C-D, S6,S7). We found that these 12.5 Hz waves were organized as TWs showing a peak at spatial frequencies around 80-90 radians/meter (spatial wavelengths around 8cm). At 12.5 Hz this corresponds to a wave velocity around 1 m/s. Note however that the energy of these TWs extended in a broader frequency range up to approx. 20 Hz.

The TWINDEX shows more specificity in terms of spatial frequency as compared to the gKFS (Figure 4 and S7) and the PO-gKFS (Figure S6). Furthermore, while gKFS and PO-gKFS suggest potential waves at the 50 Hz (line noise frequency), the TWINDEX does not detect any TWs at 50 Hz. The TWINDEX thus avoids detecting TWs when the channels show a high phase concentration, as in case of 50 Hz, by normalizing for this phase concentration. Analysis of the planar-focused TWINDEX also suggests that the spectral peak in the TWINDEX is not driven by a standing wave structure, as a clear spectral peak is found in the planar-focused TWINDEX (Figure 4 and S7).

We examined the dependence of the TWINDEX on wave direction and spatial frequency at 12.5 Hz. We find that the TWINDEX spectrum has energy at two different phases, approximately corresponding to the anterior-posterior plane, with a stronger loading on the feedback than the feedforward direction in monkey Fr (Figure 4), but a more symmetric loading in monkey Go (Figure S7). The finding that the TWINDEX shows elevated values at two opposite wave directions may either suggest that there is a standing wave structure, or could suggest substantial variability across trials. The first possibility is in principle excluded by the fact that the planar-focused TWINDEX shows a clear peak that is comparable to the regular TWINDEX. To test for the second possibility, we performed clustering of the trials based on the TWINDEX spectrum. Surprisingly, we found that the data exhibited two distinct clusters of trials. In one set of trials, TWs showed a pattern of activity going from anterior-to-posterior (with some medio-lateral component as well), while in the other set of trials, TWs showed a pattern of activity predominantly in the posterior-to-anterior direction (Figure 5 and S8). As these patterns were found in the pre-stimulus period, they were not due to trial type (e.g. standard and deviant trials).

**Figure 5:**
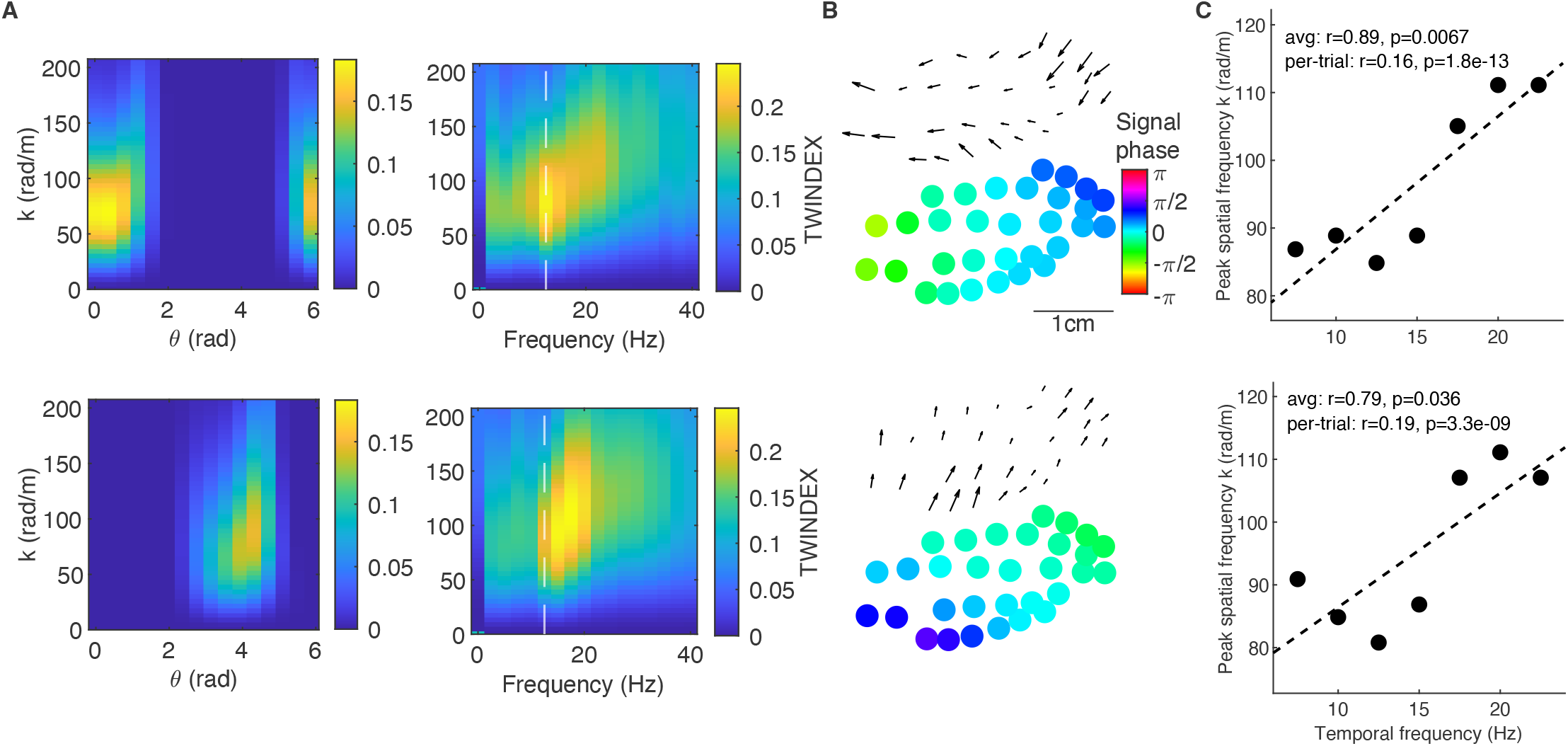
Analysis of separate clusters of trials. Trials (n=1438 trials) were clustered based on the TWINDEX as a function of spatial frequency and wave orientation, using the HDBSCAN algorithm. Top and bottom row correspond to two different clusters identified by HDBSCAN. The top row shows propagation from frontal and temporal to occipital electrodes (i.e. feedback direction; n=606 trials). The bottom row shows propagation from occipital electrodes towards frontal electrodes (i.e. feedforward direction; n=332 trials). (A): TWINDEX as a function of spatial frequency and wave direction; TWINDEX as a function of spatial and temporal frequency. (B) Phase map, obtained by aligning the phases of each trial to a circular mean of zero and then averaging across trials; gradient of the phase map. (C) Peak spatial frequency vs. temporal frequency, showing a significant positive correlation both when computing the peak spatial frequency per trial separately, or across trials.

For both anterior-to-posterior and posterior-to-anterior patterns, we found that the spatial frequency showed a positive relationship with the temporal frequency, which is consistent with the relationship 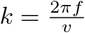, i.e., a constant propagation velocity irrespective of temporal frequency. Notably, the relationship between temporal and spatial frequency was comparable between the two groups of TW patterns.

Together, these findings suggest that alpha TWs are not exclusively related to either feedback or feedforward processing, but in a spontaneous state, rather occur in two oppositely directed propagation modes, consistent with bistable dynamics.

### Traveling waves in human EEG data

We analyzed a dataset in which 13 participants performed a visual detection task while a 64-channel EEG was recorded (Pang et al., 2020). Participants alternated 5 seconds of visual stimulation with 5 seconds of stimulus absence. The visual stimulus consisted of a white noise flicker, generating strong alpha-band oscillations in the visual cortex (VanRullen and Macdonald, 2012) (Figure 6). We analyzed both the TWs in the absence of visual stimulation (Stim-OFF, Figure 6) and those during visual stimulation (Stim-ON Figure S9). In both conditions, we observed elevated alpha power, predominantly over occipital channels. Both the gKFS and the TWINDEX spectra reveal a distinct peak in the alpha-band. Whereas the gKFS exhibited a broad peak spanning a wide range of spatial frequencies, the TWINDEX identified a well-defined peak at 20 radians/meter, as expected from its theoretical properties. This value corresponds to a propagation speed of approximately 3.2 m/s, in line with previous studies (Hughes, 1995; Patten et al., 2012). The narrow-band spatial-frequency profile suggests that alpha activity does not span a wide range of spatial frequencies, e.g., reflecting diffusive activity with a 1*/k* spectrum (Alexander and Dugué, 2024). Rather, the narrowband spatial-frequency profile suggests the presence of a genuine TW structure. This result also holds true for the planar-focused TWINDEX, obtained by discarding the standing component (Figure 6 and Figure S10). In particular, the standing-focused TWINDEX exhibits a less pronounced peak, as demonstrated by a t-test comparing the maximum values derived from the planar and standing components across the two conditions (Stim-OFF *t*(12) = 15.34, *p <* 10^−9^, 95% CI [0.0331, 0.0441]; Stim-ON *t*(12) = 17.98, *p <* 10^−10^, 95% CI [0.0338, 0.0431]). We also observed a marked difference in spatial frequency between the standing and planar waves. The planar component exhibited a higher spatial frequency, centered around 25 radians/meter, compared with approximately 18 radians/meter for the standing component, a difference confirmed by statistical testing (Stim-OFF *t*(12) = 49.52, *p <* 10^−15^, 95% CI [6.6275, 7.2375]; Stim-ON *t*(12) = 6.18, *p <* 10^−4^, 95% CI [4.2601, 8.8889]).

**Figure 6:**
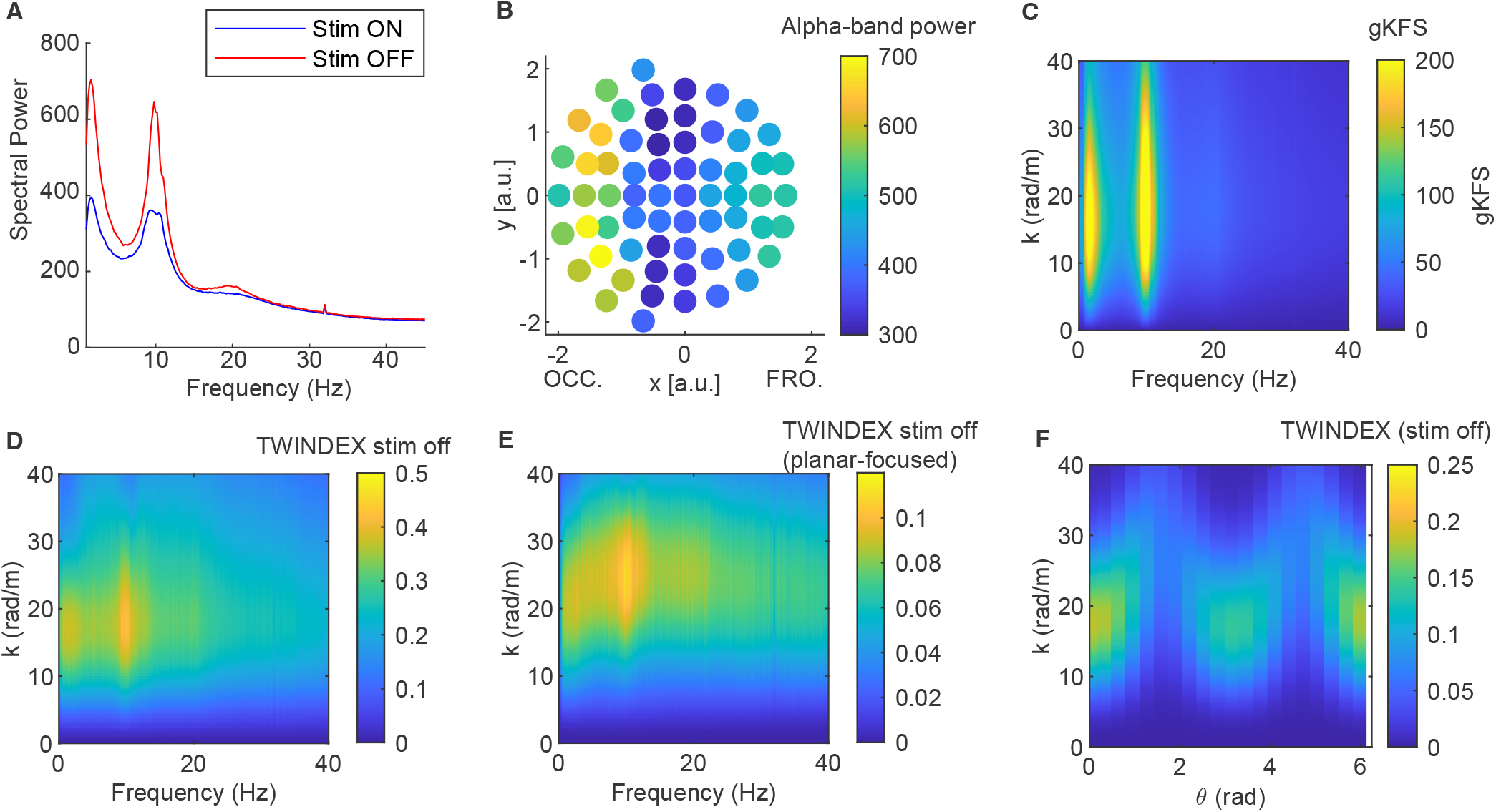
EEG results in the absence of visual stimulation. A) Averaged power distribution for all electrodes in the presence (stim ON) and absence (stim OFF) of visual stimulation. B) Topography of the average alpha-band power across participants in the Stim OFF condition. OCC. and FRO. denotes the occipital and frontal regions on the x-axis. C,D,E) Generalized *k*–*f* spectrum (gKFS), TWINDEX, and planar-focused TWINDEX applied to scalp EEG data recorded in the absence of visual stimulation. F) TWINDEX values as a function of waves’ direction and spatial frequency.

To analyze directionality, we examined the dependence of the TWINDEX on wave direction *θ* and spatial frequency *k*. This analysis shows that alpha-band TWs predominantly propagate in the frontal-to-occipital direction, corresponding to a wave direction of *θ* = 0 or equivalently 2*π* radians (Figure 6). This directional bias was particularly strong in the absence of visual stimulation, as indicated by the difference between the visual-stim on- and off-conditions (Figure 7A). Both the TWINDEX and planar-focused TWINDEX showed higher values in the alpha band in the absence of visual stimulation, with a stronger concentration of propagation directions around *θ* = 0 (2*π*) radians.

**Figure 7:**
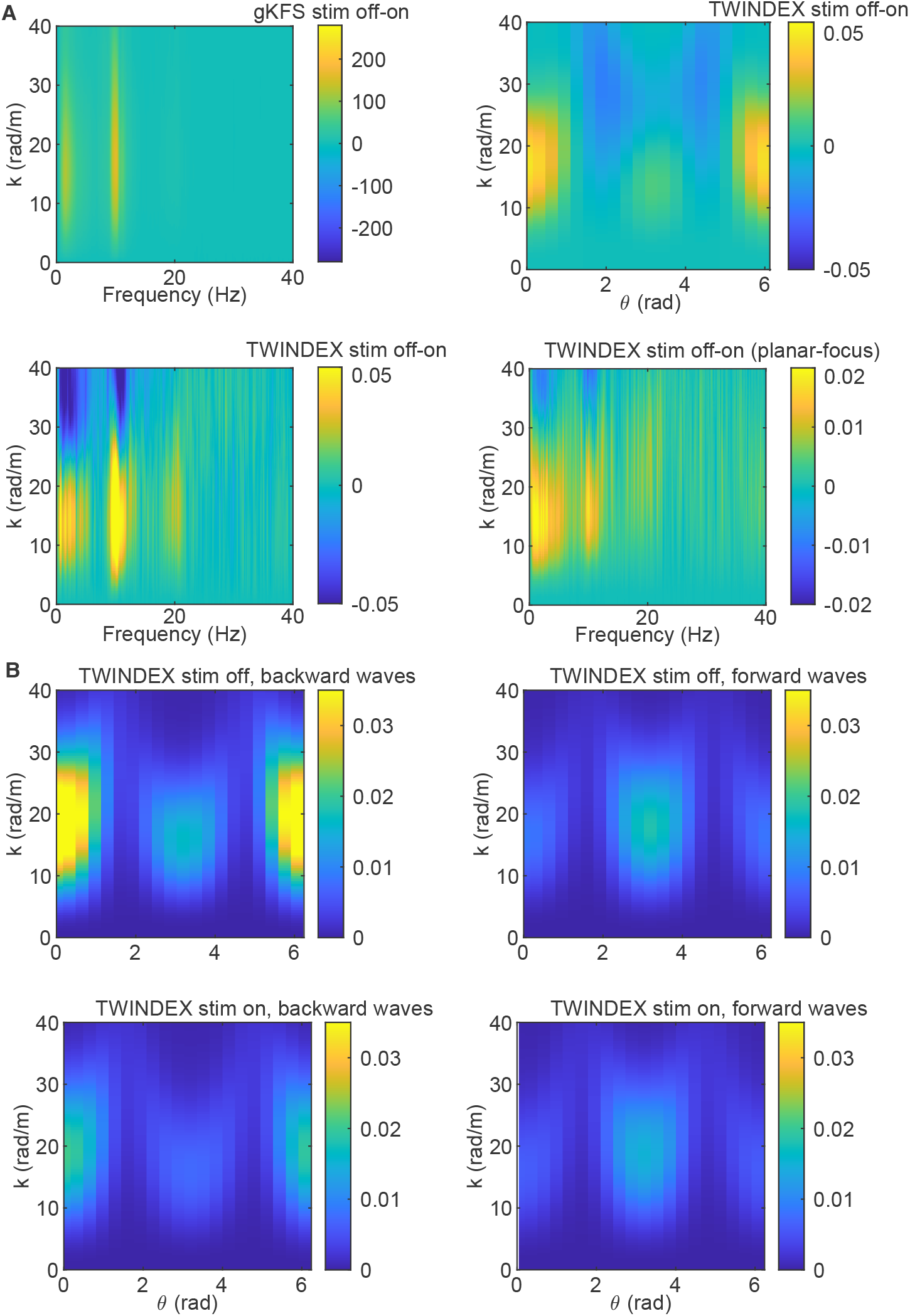
A) Difference between absence of visual stimulation (OFF) and visual stimulation (ON) for the generalized *k*–*f* spectrum (gKFS), TWINDEX as a function of the direction and of the temporal frequency, and the planar-focused TWINDEX. B) TWINDEX values as a function of waves’ direction and spatial frequency applied on data clustered based on the main propagation direction.

**Figure 8:**
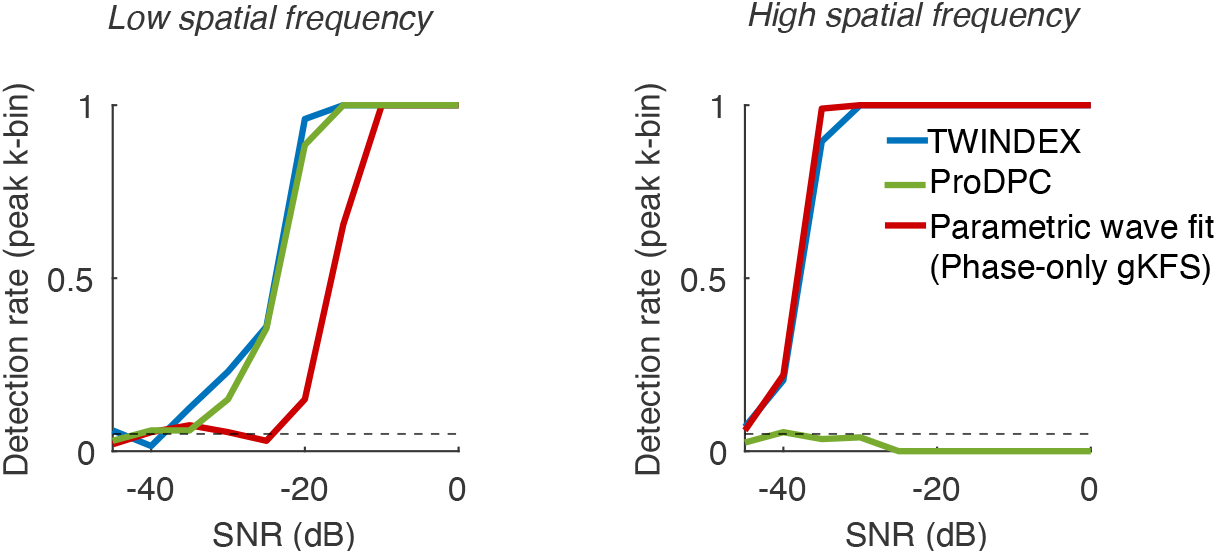
(Left) Detection rate of planar waves (significance vs. a null-distribution obtained by shuffling channels) for different methods. Green: ProDPC (projected distance-phase correlation), a proposed improved method for detection of planar waves; blue: TWINDEX; red: phase-only gKFS / planar-wave fit. Spatial frequency was low (0.5*×π* radians/m). (Right) Same as (Left), but now for a high spatial frequency, 23.5*×π* radians/m. Array size was 0.2 m.

As for the marmoset dataset, we clustered the trials according to the predominant direction of propagation, distinguishing between TWs traveling from occipital to frontal electrodes and TWs propagating in the opposite direction (Figure 7B). We then stratified 500-ms epochs according to their predominant propagation direction. Epochs with TWs propagating from occipital to frontal sensors accounted for 15% of epochs in both conditions across participants, whereas backward-propagating TWs accounted for 19% and 32% of epochs in the presence and absence of visual stimulation, respectively. Consistent with our findings in marmosets, we identified two distinct patterns of TW activity propagating in opposite directions, from occipital to frontal and from frontal to occipital sensors (Figure 7B). Together, these findings extend our observations in marmosets to human EEG, suggesting that alpha TWs are not exclusively associated with either feedback or feedforward processing, but instead occur in two oppositely directed propagation modes.

### Connection to single-trial traveling-wave detection methods

The TWINDEX can be applied in two different ways: First, to provide a spectral decomposition of TW activity by averaging across trials, which is particularly useful for data with a lower signal-to-noise ratio. Second, as we will show, the TWINDEX can be applied to detect TWs in single trials, by detecting the maximum across parameters for each trial separately. Previous methods have been proposed to detect TWs in single trials, either by fitting the parameters of a parametric wave fit (Zhang et al., 2018), or by nonparametrically correlating distance from an origin point with phases (Muller et al., 2016; Davis et al., 2020). Thus, we wondered how the TWINDEX compares with these single-trial detection methods.

In the parametric planar-wave fit (PWF) (Zhang et al., 2018), the spatial frequency and orientation of a planar wave are fitted by subtracting the predicted phases from the phase map, and maximizing the resultant vector length *R* of the residual phases (Eqs. 48 and 49). As shown in Methods (Eq. 52), the square root of the PO-gKFS evaluated at the fitted parameters *f* ^∗^, *k*^∗^, *θ*^∗^ is exactly this residual resultant vector length *R*,

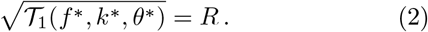

Consequently, maximizing the PO-gKFS over spatial frequency and orientation yields the same fitted parameters as PWF,

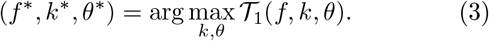

Thus, PWF can be interpreted as identifying the maximum of the PO-gKFS spectrum on each individual trial (Eq. 52).

We next consider distance-phase correlation (DPC), an alternative nonparametric single-trial measure of traveling-wave structure. The key idea of DPC is to transform 2-D space into a 1-D variable in order to apply standard circular-linear correlation coefficients. This transformation is achieved by first identifying a putative wave origin, as the point of maximum divergence in the phase gradient field (Davis et al., 2020). DPC then measures the Euclidean distance of each sensor from this origin (Eqs. 57 and 60) (Muller et al., 2016; Davis et al., 2020). The circular-linear correlation between signal phase and Euclidean distance is then computed. The DPC is designed to detect an arbitrary (non-rotating) wave form, primarily planar and radial waves (Davis et al., 2020) (i.e. no circular components). As we will show below, DPC has significant advantages compared to PWF for planar TW detection at low spatial frequencies. We further show that DPC can be further improved to address two limitations for planar wave detection: First, in its current form DPC requires to estimate a wave origin, which requires additional spatial processing such as estimation of a phase-gradient field, smoothing, or interpolation (Davis et al., 2020). Second, for a planar wave, the phase varies linearly along the true propagation axis, but not along the orthogonal direction. Yet, the Euclidean distance from the putative TW origin measures the distance across all axes, and thus DPC correlates phase with distance along axes that can be unrelated to the planar phase gradient (Figure S3).

To address these two limitations, we introduce projected distance-phase correlation (ProDPC), which is adapted specifically to planar waves (Figure S3). Rather than estimating a wave origin, we convert 2-D space into a 1-D variable by projecting the sensor locations onto a candidate orientation axis. Similar to DPC, one then computes the circular-linear correlation between phase and the resulting one-dimensional spatial coordinate (Eqs. 61 and 62). We then scan across orientations and compute the correlation per orientation (Figure S3 and S4). ProDPC measures spatial position specifically along the candidate wave-vector orientation, and avoids estimating a wave origin. For statistical testing, the orientation with the maximum ProDPC value is used.

As illustrated in Figure S3, ProDPC has a higher sensitivity than DPC in simulations of planar TWs. This finding holds true when we assume that the true wave orientation and TW origin is known, but also when these parameters need to be estimated from the data (Figure S3).

We note that the ProDPC effectively yields a spectrum that contains information about the correlation as a function of wave orientation, which allows it to directly assess whether TWs tend to cluster in a specific wave orientation across trials. However, the ProDPC can only distinguish wave orientations and not directions, in contrast to the TWINDEX, which distinguishes between waves in opposite directions (the reason is the specific circular-linear correlation coefficient, see Methods; to obtain directionality a fit of spatial frequency *k* has to be computed).

PWF and ProDPC have complementary strengths across spatial frequency regimes. In the low-*k* regime (small phase shifts close to 0 radians), DPC and ProDPC are sensitive measures of the underlying spatial relationship (Figure 8 and S4), while the PWF becomes poorly discriminative. For higher spatial frequencies and thus larger phase shifts, e.g. *> π* radians across the array, the PWF performs well, because the resultant vector length deals well with the circular nature of the predicted phase variable. By contrast, TW detection performance is reduced for DPC and ProDPC, due to the fact that the relationship between distance and the sine and cosine components of phase becomes increasingly nonlinear and non-monotonic (Figure S4). This is not a problem caused by the fact that the phases are unwrapped (the circular-linear correlation does not require unwrapping), but rather due to linearizing the phase inherent to computing the circular-linear correlation. (Note that unwrapping the phases and then computing a standard linear regression is an alternative approach, however we find that it generally leads to low TW detection performance as it tends to inflate correlations in random noise - data not shown).

The TWINDEX connects these two regimes. As shown analytically in the Methods section (Eq. 74), as *k* → 0, the TWINDEX converges approximately related to the ProDPC,

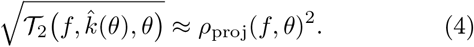

(see Figure S5). Thus, for TW detection at low *k*, TWINDEX behaves similarly to the distance-phase association captured by ProDPC. By contrast, in the high *k* regime, the TWINDEX behaves similarly to the PO-gKFS and PWF.

To test these predictions directly, we simulated planar TWs with varying signal-to-noise ratios, separately for low- and high-spatial-frequency regimes. For ProDPC, the test statistic was the maximum circular-linear correlation across orientations, and for the TWINDEX we used its maximum value across spatial frequencies and orientations. We assessed statistical significance by permuting phases across channels.

The simulation results matched the theoretical predictions. At low spatial frequencies, both the TWINDEX and ProDPC reliably detected the planar TWs, while PWF showed substantially lower sensitivity (Figure 8). At higher spatial frequencies, TWINDEX and PWF performed well, while the sensitivity of ProDPC was low (Figure 8). Together, these results show that the TWINDEX performs well in detecting planar TWs for both high and low spatial frequency regimes, in contrast to parametric wave fit and nonparametric distance-phase correlation methods.

## Discussion

We proposed the TWINDEX (Traveling Wave Index), which provides a full and improved spatio-temporal spectral characterization of neural TWs. We showed through simulations that the TWINDEX 1) is suited for irregularly sampled and non-rectangular grids; 2) is well behaved for both low and high spatial frequencies; 3) provides interpretable measures of wave strength; and 4) can distinguish genuine (unidirectional) planar waves from standing waves. Furthermore, we prove analytical relations to classic parametric planar-wave fits and distance-phase correlations (Eqs. 52 and 74), and show that the TWINDEX combines the strength of both methods, while adding a full spatio-temporal spectral characterization. We apply these methods to discover spatio-temporal TWs in human EEG and marmoset ECoG data. In marmosets, we show the existence of alpha/low-beta 12.5 Hz global planar waves traveling around 1 m/s. These spontaneous waves cluster in two types across trials, going either in the feedforward or in the feedback direction (with some propagation in medio-lateral plane), revealing two dominant, oppositely directed propagation modes across trials, consistent with bistable dynamics, with the feedback type being the most common type. In humans, we show the existence of alpha waves that travel at physiologically plausible wave speeds around 3 m/s, and are not explained as standing waves. These waves predominantly travel in the feedback direction, especially in the absence of visual stimulation.

### Comparison to 2D-FFT

The proposed gKFS has significant advantages compared to the 2D-FFT approach, which was developed by (Alamia and VanRullen, 2019) to quantify TWs in human scalp-EEG recordings, specifically along midline electrodes. The 2D-FFT method has since been applied to examine the functional role of alpha-band TWs across multiple cognitive domains. These include visual attention (Alamia et al., 2023), working memory (Zeng et al., 2024; Luo and Ester, 2024) and altered states of consciousness (Bailey et al., 2025; Alamia et al., 2020). We repeated some of the analyses (feedforward-feedback for stim-on and stim-off) in those papers and could replicate the previous conclusions using the TWINDEX. The methods here go beyond the 2D-FFT in several ways: First, the gKFS and TWINDEX extends the decomposition in two spatial dimensions, apply them to irregular and non-rectangular arrays, and are appropriate for low spatial frequencies (while the minimum frequency that the 2-D FFT resolves is already relatively high as it spans one cycle). Second, the TWINDEX then applies appropriate scaling and normalization to yield a dimensionless, interpretable measure of wave strength that is not artificially inflated at low spatial frequencies (i.e. small phase shifts) and is not biased by signal power. Compared to an amplitude-normalized gKFS, the TWINDEX yields superior wave detection at lower spatial frequencies and a much larger dynamic range. Third, we show that by examining the asymmetry of the TWINDEX as a function of wave direction, the TWINDEX can be weighted in order to be selectively focused on either planar or standing waves. This is crucial as standing waves should be a common phenomenon in electrophysiological data caused by dipoles (Pesaran et al., 2018). Fourth, we can analytically link the values of the TWINDEX to interpretable single-trial estimates of wave strength, namely the parametric planar-wave fit and distance-phase correlation (Eqs. 52 and 74).

There is also a relation of the gKFS with the singular value decomposition (SVD) based method. For spatially broadband spectra, the SVD method can yield similar results to Fourier Decomposition, as it will decompose the spectra in orthogonal components (Alexander and Dugué, 2024). Like SVD, overlapping waves can be extracted using TWINDEX and PO-gKFS, as their spectral representation can reveal multiple distinct peaks or components. SVD may share some of the same limitations we discussed for the gKFS, if we take it as a measure of wave strength. Furthermore, the SVD may converge on frequencies that equal the discrete frequencies allowed by the Discrete Fourier Transform, as those constitute orthogonal base functions (Alexander and Dugué, 2024). This means that the TWINDEX approach may have advantages for the low spatial frequency regime, which we argued is typical for brain data.

### Comparison to single-trial detection techniques

The parametric planar-wave fit method (PWF) was introduced by (Zhang et al., 2018). Here the signal phase for each sensor is predicted from the spatial positions on the array relative to some point of origin, weighted by estimated regression coefficients. A coefficient of explained variance is computed as the resultant vector length. We showed here that taking the maximum PO-gKFS yields a computationally efficient method to obtain the PWF (Eq. 55). The TWINDEX moves beyond the PWF by providing an adequate measure of TW strength for lower spatial frequencies, filtering out standing waves, and computing a full spatio-temporal spectral decomposition that can be used to pool evidence across trials.

Distance-phase-correlation (DPC) is a TW detection method for single trials based on performing circular-linear regression between phase and a 1-D distance to some point of origin (Muller et al., 2018, 2014, 2016; Davis et al., 2020). Using DPC, planar wave fits can be compared with radial wave and rotating wave fits for single trials (Muller et al., 2016).

We developed here, for planar waves, an improvement to the distance-phase correlation (DPC), which we call the projected distance-phase correlation (ProDPC). ProDPC bypasses the need to find a wave origin and measuring distances relative to that origin, which is problematic for planar waves. ProDPC effectively beam-scans in all wave orientations, projecting the spatial positions onto these wave orientations, and then computing phase-distance correlations. The ProDPC measure can be used for single-trial detection of waves, but can also be averaged across trials to examine the modulation of the ProDPC as a function of temporal frequency and wave direction. The TWINDEX moves beyond DPC/ProDPC by providing a measure that behaves well for higher spatial frequencies (phase shifts larger than *π* radians), and that can filter out standing waves.

### TW detection and spatial frequencies

The relation 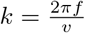 can be used to assess which spatial frequency regimes are typical for brain data. If information travels via multi-synaptic chains over unmyelinated axons, speeds around 0.5 m/s may be typical (Muller et al., 2014), while communication over myelinated axons can have axonal conduction speeds between 1-10 m/s. However one must consider that the measured propagation over multiple areas may be considerably slower, given that one has to factor in the rise time of excitatory post-synaptic potentials and processing within each area, which involves multiple synapses (e.g. flow from cortical Layer 4 to 2/3) (Dugué and Chavane, 2025). Communication over regions is usually associated with temporal frequencies between 1-100 Hz (delta, theta, alpha, beta, gamma). Higher spatial frequencies would occur e.g. if information propagates via higher gamma frequencies, while lower spatial frequencies would occur if information propagates via lower temporal frequencies (Vinck et al., 2025). Sensor arrays may be on the order of mm’s to tens of cm’s, depending on whether TWs are quantified at a meso- or macroscopic scale and the size of the brain (e.g. distance of 1-2 cm between visual areas in macaque and human, but mm in mice and marmoset). Considering these numbers, lower spatial frequencies (i.e. phase shifts across the array smaller than 2*π* radians) of neural TWs can easily occur under some conditions (smaller brain, higher propagation speeds, lower temporal frequencies). Higher spatial frequencies may be expected for transmission at higher temporal frequencies (e.g., feed-forward) and larger brain size (macaque, human). However, higher spatial frequencies and phase shifts can be caused by other noise factors or e.g. diffuse activity that covers a broad range of spatial frequencies (Alexander and Dugué, 2024).

The fact that neural TWs may typically have relatively low spatial frequency contents is an important limitation for decomposition techniques whose first resolvable frequency corresponds to effectively 2*π* radian phase shifts (2-D FFT, SVD) (Alamia and VanRullen, 2019; Alexander and Dugué, 2024) (similar to a 1-D FFT). Furthermore, we showed distinct advantages of quantification using DPC and TWINDEX for lower spatial frequencies, compared to PWF.

### Beyond planar and standing waves

The main limitation of the present work is that the spatiotemporal decomposition of TWs proposed here is based on spatial Fourier base functions, i.e. planar waves. This decomposition can distinguish planar and standing waves, and can in principle distinguish overlapping components with different temporal and spatial frequencies. Thus, we argue that the TWINDEX is in some sense the ideal method to characterize planar waves, and have summarized its advantages above. However, the assumption is that the TW can be decomposed into sums of planar waves. For radial waves that behave as *ϕ* = *k***r** where *r* is the Euclidean distance to a point of origin, a decomposition in terms of planar waves is not well suited. In such a case, the TWINDEX will yield values close to 0, as fitting a planar wave in any direction will not reduce the dispersion of phases. The PO-gKFS and gKFS will yield energy across all wave directions with a broad lobe towards *k* = 0. Thus, for radial waves or rotating TWs, parametric or nonparametric fits of these TWs need to be computed.

### Empirical findings

Previous studies have applied multiple approaches to quantify alpha-band TWs in human scalp-EEG (Ito et al., 2005; Lozano-Soldevilla and VanRullen, 2019; Nunez et al., 2001; Alamia and VanRullen, 2019; Schwenk and Alamia, 2026), yet interpretation of these findings has been constrained by limitations inherent to EEG measurements (Nunez and Srinivasan, 2006, 2014; Grabot et al., 2025).

The method introduced here improves the quantification of alpha-band dynamics in EEG recordings by mitigating the influence of volume conduction, which induces coherent phase relationships across electrodes and can artificially inflate estimates of gKFS and PO-gKFS. The TWINDEX is not artificially inflated by coherent in-phase activity across channels, as such a pattern will result in a TWINDEX of zero (due to the normalization of the TWINDEX). As an example, the TWINDEX completely suppressed the line noise in the EEG and marmoset recordings, in contrast to the gKFS and PO-gKFS. The planar-focused TWINDEX approach further addresses the problem of standing waves generated by volume-conducted single dipolar sources. In the empirical data we found robust spectral peaks in the planar-focused TWINDEX.

Our results support the hypothesis that in humans during rest, spontaneous alpha TWs are biased towards the feedback direction (Alamia et al., 2023; Samaha et al., 2015), independent of cognitive functions (Doesburg et al., 2016; Mayer et al., 2015; Sauseng et al., 2005). The tendency to generate these TWs may depend on the configuration of anatomical connections which, combined with cortical delays, produce these waves in the feedback direction (Budzinski et al., 2023). The observation of TW speeds around 3 m/s indeed suggest potential propagation via myelinated axons. However, it remains possible that observed alpha TWs reflect a more anterior source at an earlier alpha phase that is phase-coherent with a more posterior source at a later alpha phase (Zhigalov and Jensen, 2023; Orsher et al., 2024). Furthermore, the observed feedback pattern reflects the distribution of power across the cortex and arises as a consequence of communication, rather than a mechanism for communication (Schneider et al., 2021; Vinck et al., 2025, 2023; Pesaran et al., 2018; Spyropoulos et al., 2024; Schneider et al., 2023).

We showed the existence of spontaneous global planar-like TWs in marmoset ECoG signals around 12 Hz. Trials clustered in two wave patterns in opposite directions, one going in the feedforward and one in the feedback direction, such that the TWINDEX contained peaks at two different wave directions. This indicates a potentially bistable system that takes either one of two configurations, perhaps depending on e.g. the excitability or initial state in that particular trial, relying on the same eigenmode (Budzinski et al., 2023). Spontaneous TWs in a similar frequency range have previously been observed in awake marmosets, with spectral content extending from approximately 10–30 Hz in area MT (Davis et al., 2020). Our results extend these observations by identifying a spectrally localized traveling-wave component around 12.5 Hz across large-scale marmoset ECoG recordings.

The findings here offer a different perspective on the spectral organization of TWs than in (Alexander and Dugué, 2024), who described a broad 1*/k* spectrum of energy in EEG data. Rather, we find that energy is focused in a limited band of spatial frequencies, that are considerably lower than the first spatial frequency that can be resolved through singular value decomposition or classic Fourier decomposition techniques. This does not indicate diffusive activity resulting in a broadband 1*/k* spectrum, but is rather consistent with mechanisms of interacting oscillators, e.g. as in (Budzinski et al., 2023).

### Outlook

In sum, we provide a new set of techniques for a full spatiotemporal decomposition of neural data, to detect neural TWs, with interpretable measures of wave strength. We showed in empirical data that this technique sheds new light on the large-scale organization of alpha/beta waves in cortex, showing large-scale alpha/beta waves that are localized in both spatial and temporal frequency as well as wave direction, and which occur in two oppositely directed propagation modes across trials, consistent with bistable dynamics. TWs are increasingly recognized as a fundamental organizing principle of brain activity, coordinating neural processing across space and time (Cruddas et al., 2025; Muller et al., 2018; Sato et al., 2012; Ermentrout and Kleinfeld, 2001), and linking micro- to meso- to macro-scales. We expect that the proposed TWINDEX method will provide new avenues for exciting discoveries on the relation between TWs and cognition and perception.

## 2. Methods

To define the generalized *k*–*f* spectrum (gKFS), we first consider the 3-D Discrete Fourier Transform (DFT). Because the multidimensional DFT is separable, the temporal and spatial transforms can be computed sequentially (Oppenheim and Schafer, 2010). We first Fourier-transform the signals in time, yielding the complex-valued temporal spectrum 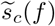 for each channel *c*, and then compute the spatial Fourier representation.

Although we use the temporal DFT here for concreteness, the same spatial analysis can be applied to complex-valued frequency-domain signals obtained from wavelets or band-pass filtering followed by the Hilbert transform.

For a rectangular, regularly sampled sensor grid, let *x*_j_ and *y*_*ℓ*_ denote the physical sensor coordinates along the two spatial dimensions. The spatial Fourier transform at temporal frequency *f* can then be written as

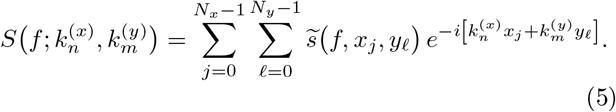

For regularly spaced samples, the discrete angular spatial frequencies are

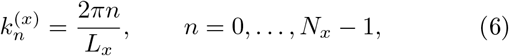

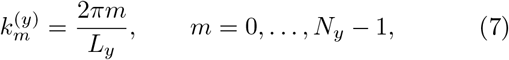

where *L*_x_ and *L*_y_ denote the spatial periods associated with the sampled grid.

We next reparameterize the two-dimensional spatial-frequency vector **k** = (*k*^(x)^, *k*^(y)^) in polar coordinates,

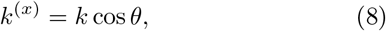

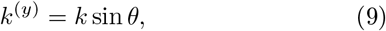

where *k* ≥ 0 denotes the angular spatial frequency (wavenumber), with units rad/m, and *θ* denotes the orientation of the spatial wave vector. For each orientation *θ*, we define the projected spatial coordinate of channel *c* as

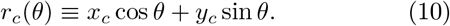

The spatial spectrum at temporal frequency *f* can then be written as

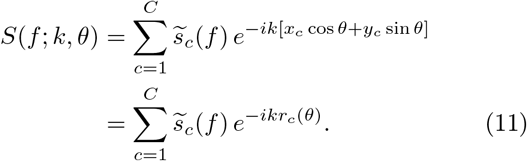

Here, (*x*_c_, *y*_c_) denotes the physical location of channel *c*, and *C* is the total number of channels.

By writing the transform as a sum over sensor locations, we remove the requirement that sensors lie on a regularly sampled rectangular grid. This formulation therefore applies equally to regular and irregular sensor layouts. Moreover, unlike the standard DFT, *k* need not be restricted to the discrete Fourier frequencies 2*πn/L*. Instead, Eq. 11 can be evaluated for an arbitrary, and potentially densely sampled, set of spatial frequencies *k* and directions *θ*. This allows spatial frequencies below the conventional Fourier-bin spacing to be evaluated, although the analysis resolution remains limited by the spatial aperture and geometry of the sensor array. For a regularly sampled grid with sensor spacing Δ, the angular spatial Nyquist frequency is

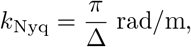

corresponding to a minimum resolvable wavelength of 2Δ. For spatial frequencies above this limit, spatial aliasing will occur.

Computing the spectrum in this way is closely related to beam scanning for planar waves with different spatial frequencies and directions (Van Trees, 2002; Johnson and Dudgeon, 1993; Devaney, 1981; Balanis, 2016; Rost and Thomas, 2002). Specifically, we define the planar-wave Fourier kernel at sensor *c* as

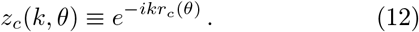

Collecting these values across channels into the vector **z**(*k, θ*), the spatial spectrum can be written compactly as

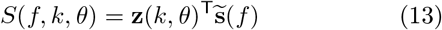

We define the generalized *k*–*f* spectrum (gKFS) as

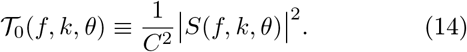

The normalization by *C*^2^ removes the dependence of the spectrum on the number of channels. The gKFS nevertheless remains amplitude-sensitive, as channels and frequencies with larger spectral amplitudes contribute more strongly to *T*_0_.

### Phase-only gKFS

The gKFS defined above is sensitive to spectral amplitude, such that channels and frequencies with larger amplitudes contribute more strongly to the spectrum. In many applications, however, we are interested specifically in the spatial phase relationships across channels, independently of their amplitudes. We therefore define a phase-only version of the gKFS.

We first decompose the complex-valued signal coefficient at each channel as

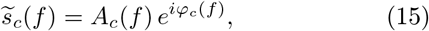

where 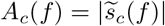 and *φ*_c_(*f*) denote its amplitude and phase, respectively. We remove the amplitude information by normalizing each coefficient to unit magnitude,

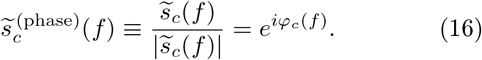

For a candidate planar wave with spatial frequency *k* and direction *θ*, we define the residual phase at channel *c* as

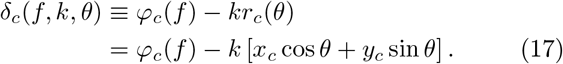

The phase-only spatial match is then given by the mean resultant vector of these residual phases,

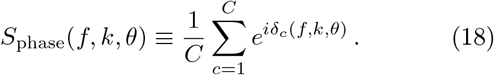

We define the phase-only gKFS (PO-gKFS) as the squared magnitude of this mean resultant vector,

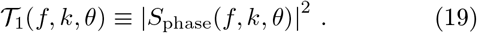

Because each channel contributes a unit-magnitude phasor, 0 ≤ *T*_1_ ≤ 1. A value of *T*_1_ = 1 indicates perfect phase alignment with the candidate planar wave, whereas smaller values indicate increasing dispersion of the residual phases. We use the squared resultant length so that *T*_1_ has the same quadratic form as the amplitude-sensitive spectrum *T*_0_.

This construction is closely related to phase-only beam-forming in array processing (Mecklenbräuker et al., 2020; Fridman, 2005). It is also mathematically related to phase-locking measures such as the phase-locking value (PLV), which quantify phase consistency by averaging unit-magnitude phasors (Lachaux et al., 1999; Pikovsky et al., 2001; Vinck et al., 2010). Here, however, the phasors are averaged across sensors after subtracting the spatial phase pattern predicted by a candidate planar wave.

### The TWINDEX

Although the PO-gKFS provides a normalized measure of phase alignment with a candidate planar wave, it becomes poorly discriminative at low spatial frequencies. The reason is that, when the spatial phase gradient across the sensor array is small, the phases are already strongly concentrated before any spatial steering is applied.

Recall that the phase-only spatial match is

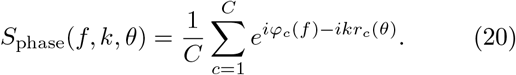

At *k* = 0, the spatial steering term vanishes, such that

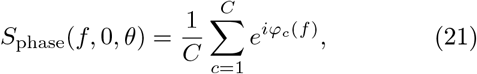

which is independent of *θ*. Consequently,

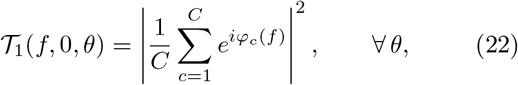

which measures only the overall phase concentration across channels and contains no information about wave direction.

This becomes particularly problematic when the true spatial frequency is low. Consider the planar-wave phase model

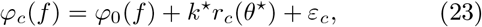

where *k*^⋆^ and *θ*^⋆^ denote the true spatial frequency and orientation, and *ε*_c_ denotes phase noise. If the phase gradient across the sensor array is small, such that

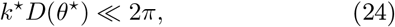

where

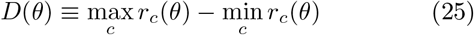

is the projected aperture of the sensor array, then the phases differ only weakly across channels. As a result, the resultant vector length is close to one, and thus *T*_1_(*f*, 0, *θ*) →1 even though it does not depend on *θ* and does not identify any wave direction. More generally, when *kD* ≪ 2*π*, steering vectors corresponding to nearby values of *k* and *θ* differ only weakly across the sensor array, producing a broad and weakly discriminative low-*k* component in the PO-gKFS.

To obtain a measure of wave strength that remains interpretable in this regime, we quantify how much a candidate planar wave reduces the circular dispersion of the observed phases relative to their original dispersion. We first define the resultant vector length and circular variance of the original phases as

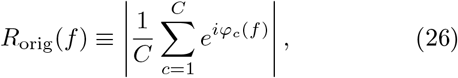

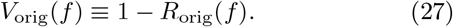

For a candidate wave (*k, θ*), we similarly define the resultant vector length of the residual phases as

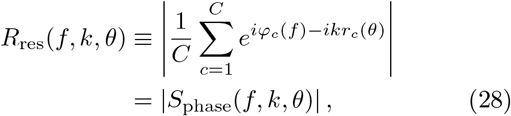

and the corresponding residual circular variance as

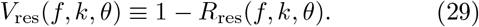

Note that

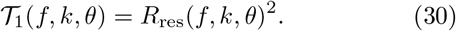

We then define the relative reduction in circular variance produced by the candidate wave as

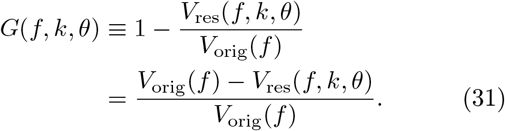

This quantity is analogous to a coefficient of determination: *G >* 0 indicates that removing the candidate planar-wave phase gradient reduces circular variance, whereas *G <* 0 indicates that it increases circular variance.

We define the Traveling Wave Index (TWINDEX) as the squared non-negative fractional reduction in circular variance,

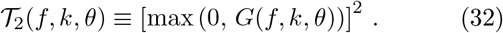

When *V*_orig_(*f*) = 0, no spatial phase variance is available to be explained, and we define *T*_2_(*f, k, θ*) = 0. In practice, to avoid numerical instability when the original phase variance is very small, we can add a very small positive constant *ε* to the denominator and define the variance-reduction coefficient as

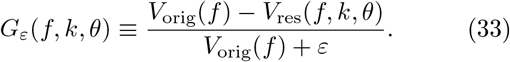

When the original phases are broadly distributed such that *V*_orig_ ≈ 1, we have *G* ≈ *R*_res_, and therefore

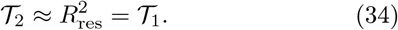

By construction, TWINDEX removes the uninformative *k* = 0 component. At *k* = 0,

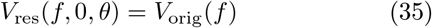

for all *θ*, and therefore

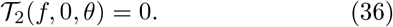

Thus, TWINDEX measures the improvement in spatial phase alignment produced by a candidate wave relative to the phase concentration already present in the unsteered data, rather than phase concentration itself.

For analyses across *P* trials, we first compute the variance-reduction coefficient separately for each trial,

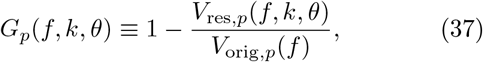

and define the across-trial TWINDEX as

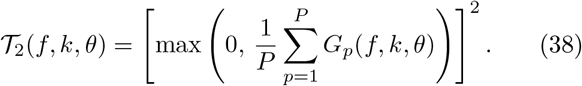

Applying the non-negativity constraint only after averaging avoids introducing an upward bias by separately truncating negative values on individual trials.

### Angular structure of TWINDEX: planar, standing, and radial waves

The angular structure of the TWINDEX spectrum provides information about the geometry of the underlying wave pattern. A planar traveling wave is expected to produce a dominant peak at a single direction, whereas a standing wave produces approximately opposing peaks separated by *π*.

For each temporal frequency *f* and spatial frequency *k*, let

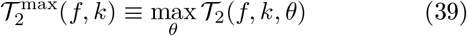

denote the maximal TWINDEX across directions. We normalize the angular spectrum as

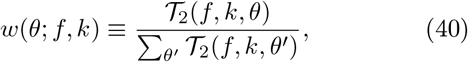

such that

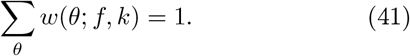

We characterize the angular distribution using its first and second circular moments. The magnitude of the first circular moment is

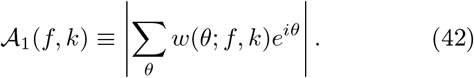

*A*_1_ measures directional asymmetry of the angular spectrum. A value close to one indicates that the spectrum is concentrated around a single direction, as expected for a planar traveling wave. By contrast, *A*_1_ is small for angular distributions that are symmetric under opposite directions, including both standing-wave and approximately isotropic patterns.

The magnitude of the second circular moment is

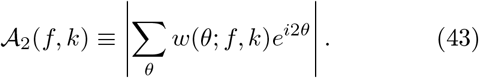

*A*_2_ measures axial structure of the angular spectrum. It is large when the spectrum is concentrated around a common direction axis, including a pair of opposing peaks at *θ* and *θ* + *π*, as expected for a standing wave. An approximately uniform angular distribution yields small values of both *A*_1_ and *A*_2_. These quantities should be computed per trial separately, and this we omit the dependence of trial indices in the following definitions:

#### Planar-focused TWINDEX

We define the planar-focused TWINDEX as

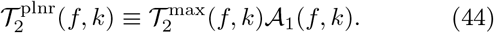

This measure emphasizes spectra dominated by a single wave direction while retaining the overall magnitude of the TWINDEX spectrum.

#### Standing-focused TWINDEX

A standing wave is characterized by strong axial structure but little directional asymmetry. We therefore define the standing-focused TWINDEX as

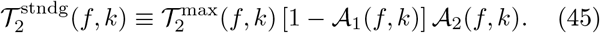

The factor 1 − *A*_1_ suppresses strongly unidirectional planar-wave patterns, whereas *A*_2_ is selective for angular spectra with an opposing two-lobed structure.

### Relation to the parametric planar-wave fit

We next derive the relationship between the PO-gKFS and the parametric planar-wave fit (PWF). In PWF, the spatial phases on a single trial are modeled by a planar phase gradient,

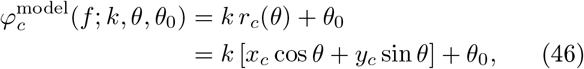

where *k, θ*, and *θ*_0_ denote the spatial frequency, wave-vector direction, and global phase offset, respectively. The wave parameters are typically estimated using a grid search or related optimization procedure (Zhang et al., 2018).

For any candidate spatial frequency and direction (*k, θ*), we define the phase residuals as

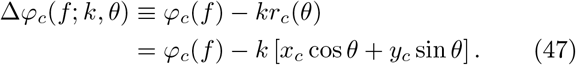

The corresponding phase-alignment objective is the resultant vector length

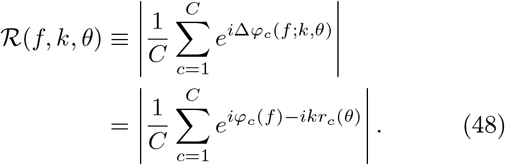

The global phase offset *θ*_0_ does not affect the resultant vector length, because subtracting a common phase from all residuals rotates the resultant vector without changing its magnitude. It therefore does not need to be explicitly optimized when estimating *k* and *θ*.

PWF estimates the wave parameters by maximizing this objective,

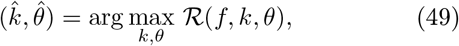

and the residual vector length of the fitted wave is

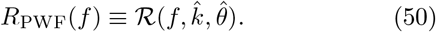

The PWF and the PO-gKFS therefore yield identical estimates of the optimal spatial frequency and direction,

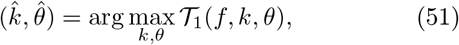

and the goodness-of-fit of the fitted PWF model satisfies

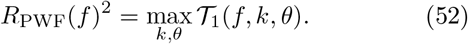

Thus, PWF can be interpreted as identifying the maximum of the PO-gKFS on each trial, whereas the PO-gKFS retains the full dependence of phase alignment on spatial frequency and direction.

This equivalence also shows that the low-spatial-frequency limitation described above is not specific to the PO-gKFS. Because residual-vector-length PWF optimizes the same phase-alignment criterion, it likewise becomes poorly discriminative when the spatial phase gradient across the sensor array is small. TWINDEX addresses this limitation by quantifying the reduction in circular variance relative to the phase dispersion present before spatial steering.

The spectral formulation also provides a convenient vectorized implementation of PWF. Let the *C* × *M* matrix **Z** contain the planar-wave kernels for *M* tested combinations of spatial frequency and wave direction,

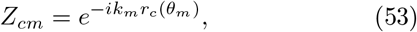

and let the *F* × *C* phase-only data matrix be defined as

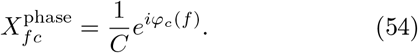

The matrix product

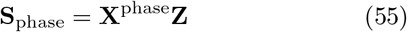

then contains *S*_phase_(*f, k*_m_, *θ*_m_) for all temporal frequencies and candidate wave parameters simultaneously. This allows the complete parameter search to be implemented using a single matrix multiplication rather than explicit loops over spatial frequencies and directions.

Although PWF is typically applied separately to individual trials, the spectral formulation also allows wave parameters to be estimated from a collection of trials when a common propagation mode is of interest. In this case, the PO-gKFS is first averaged across *P* trials and the peak of the resulting spectrum is determined,

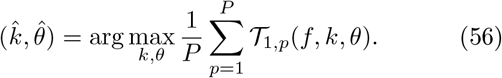

This estimates the spatial-frequency and wave direction combination showing the strongest phase alignment on average across trials.

### Distance-phase correlation and projected distance-phase correlation

A previously introduced approach for detecting traveling waves quantifies the relationship between signal phase and spatial distance (Muller et al., 2014; Davis et al., 2020; Muller et al., 2016). We refer to this class of methods as distance-phase correlation (DPC).

In DPC, a one-dimensional spatial coordinate is first constructed by estimating a reference point (*x*_0_, *y*_0_) associated with the origin of the wave, for example from the divergence of the phase-gradient field (Davis et al., 2020) or from the location of minimum phase latency (Muller et al., 2014). The Euclidean distance of each channel from this reference point is then computed as

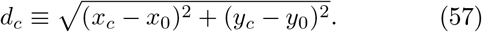

The association between distance *d*_c_ and signal phase *φ*_c_(*f*) is quantified using a circular-linear correlation.

Specifically, we define

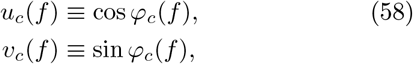

and let

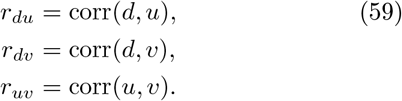

The standard squared circular-linear correlation coefficient is then

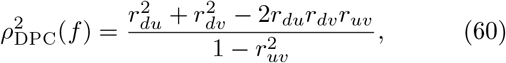

with *ρ*_DPC_(*f*) ∈ [0, 1]. Unlike the gKFS and TWINDEX, DPC yields a single measure of phase–space association rather than a spectrum over spatial frequencies.

DPC can be used to detect arbitrary, non-rotating TWs, including radial and planar waves. For planar traveling waves, however, the use of Euclidean distance from an estimated origin has several disadvantages. First, estimating the origin can be unreliable when the signal-to-noise ratio is low. Second, approaches based on the phase-gradient field and its divergence generally require a regularly sampled spatial grid, or interpolation onto such a grid. Third, Euclidean distance is not the natural spatial coordinate for a planar wave. For a planar wave, phase varies along the wave-vector direction but is constant along the orthogonal direction. Consequently, changes in Euclidean distance caused by displacements orthogonal to the wave direction introduce spatial variation that is unrelated to phase and can attenuate the distance–phase association (Figure S3).

We therefore define a projected distance-phase correlation (ProDPC) that avoids the need to estimate a wave origin. Rather than computing Euclidean distance from a reference point, we project each sensor location onto a candidate orientation *θ*. Using the projected spatial coordinate defined above,

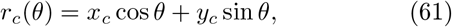

we compute the circular-linear correlation between *r*_c_(*θ*) and *φ*_c_(*f*) for each candidate orientation *θ*. Specifically, the ProDPC coefficient 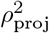 (*f, θ*) is obtained from Eq. 60 after replacing *d*_c_ by *r*_c_(*θ*).

For TW detection, we can then find the maximum circular-linear correlation across orientations,

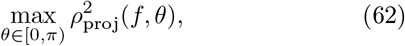

ProDPC removes the need to identify a wave origin, applies directly to irregular and non-rectangular sensor layouts, and uses the spatial coordinate naturally associated with a planar phase gradient. Unlike the gKFS and TWINDEX, however, it does not estimate spatial frequency *k* and therefore does not yield a spatial-frequency spectrum.

### Analytic relation between TWINDEX and ProDPC at low spatial frequency

We next derive the relationship between TWINDEX and ProDPC in the low-spatial-frequency regime. We assume that the data are generated by a planar wave with spatial frequency *k*^⋆^ and orientation *θ*^⋆^, such that

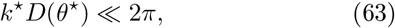

and that the phases are sufficiently concentrated for phase wrapping to be negligible. We further consider candidate directions *θ* close to the true wave direction and optimize TWINDEX over spatial frequency *k* for each direction.

For a candidate orientation *θ*, we center the projected spatial coordinate,

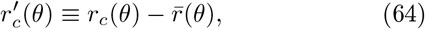

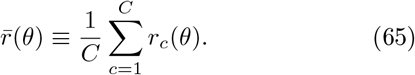

We similarly define the mean phase direction

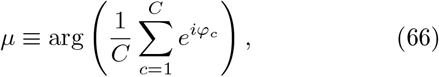

and the phase deviations

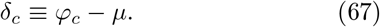

For concentrated phases, |*δ*_c_ | ≪ 2*π*, such that the circular variance can be approximated by half the ordinary variance,

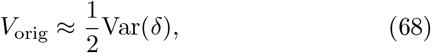

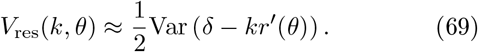

Recalling the variance-reduction coefficient

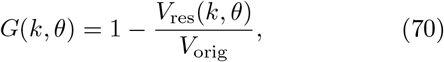

we therefore obtain

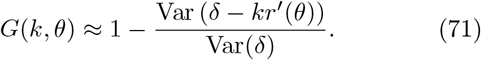

For a fixed orientation *θ*, the residual variance is minimized by

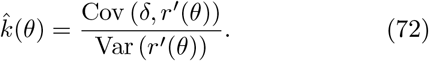

Substituting this value gives

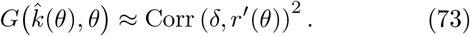

Since *T*_2_ = [max(0, *G*)]^2^, and *G >* 0 around the fitted wave,

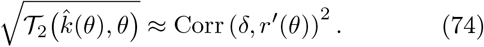

We next show that ProDPC reduces to the same quantity in the low-spatial-frequency regime. For a given orientation *θ*, ProDPC computes the circular-linear correlation between *r*_c_(*θ*) and *φ*_c_. Because the circular-linear correlation is invariant to a common rotation of all phases, we can equivalently use

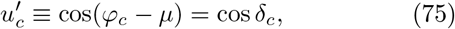

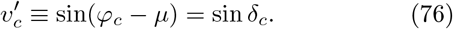

For concentrated phases,

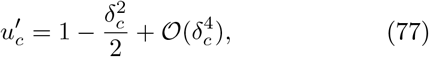

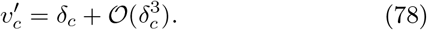

Thus, *v*^′^ captures the leading linear relationship between phase and projected position, whereas *u*^′^ captures second-order variation in the phase deviations. For a locally linear planar phase gradient, the first-order term 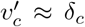 captures the signed relationship between phase and projected position, whereas the second-order term 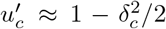 primarily reflects the magnitude of the phase deviation rather than its sign. For approximately symmetric spatial sampling and phase fluctuations, this quadratic term contributes little additional information about projected position. In this regime, the circular-linear coefficient of determination therefore reduces approximately to

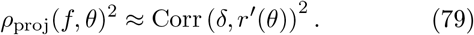

Thus, in the low-spatial-frequency regime, the variance-reduction coefficient underlying TWINDEX approaches the coefficient of determination of ProDPC (Eq. 74). By contrast, when the original phases are broadly distributed, *V*_orig_ ≈ 1, TWINDEX approaches the PO-gKFS *T*_1_ (Eq. 34). TWINDEX therefore connects the low-*k* regime captured by ProDPC with the regime in which the phase-only spectral measure is well behaved.

### Scalp EEG recordings

EEG data were acquired from 13 healthy participants during a visual detection task (Pang et al., 2020). Each participant completed five sessions, each comprising 10 experimental blocks of six trials. Trials consisted of 5 s of visual stimulation (stimulus ON), followed by 5 s of a blank screen (stimulus OFF). Throughout each trial, participants maintained central fixation and covertly attended to the stimulus in order to detect a brief luminance-decrement target.

The data were preprocessed as in the original publication (Pang et al., 2020). Continuous EEG was recorded at 1024 Hz using a 64-channel active BioSemi system, with three additional electrodes for ocular activity monitoring. Preprocessing was conducted in EEGLAB using custom scripts and included both target-present and target-absent trials. Noisy channels were removed, data were downsampled to 160 Hz, and line noise was attenuated using a 47–53 Hz notch filter. Signals were re-referenced to the common average, and slow drifts were eliminated with a high-pass filter (cutoff >1 Hz). Epochs containing eye movements, blinks, muscle artifacts, or other contamination were rejected from further analysis.

### Marmoset ECoG recordings

ECoG data were collected from two marmosets (FR [32 electrodes] and GO [64 electrodes]) during a roving odd-ball paradigm (Canales-Johnson et al., 2021; Gelens et al., 2024). In this task, sequences of identical tones (3, 5, or 11 repetitions) were presented continuously. Within each sequence, the first tone functioned as an unexpected deviant, whereas the final tone served as an expected standard. Pure tones of varying frequencies were delivered with fixed duration and interstimulus intervals. To equate the number of standard and deviant trials, only the final standards were included in the analysis.

ECoG signals were recorded at 1 kHz using a multi-channel acquisition system (0.3–500 Hz bandpass), rereferenced to the common average, high-pass filtered at 0.5 Hz, and segmented into pre-stimulus epochs of 400 ms. This yielded 1440 standard and 1440 deviant epochs for GO, and 720 standard and 720 deviant epochs for FR.

Fourier Transforms were computed with a Hanning taper. To separate the trials in clusters, we used the HDBSCAN algorithm, and computed the Euclidean distance between TWINDEX patterns across trials as the input to the HDBSCAN algorithm.

## Author contributions

Conceptualization of methods: MV. Analysis marmoset data: MV, JS, ACJ, MK. Analysis human EEG data: AA, JS. Mathematical analysis: MV. Numerical simulations: MV. First manuscript draft: MV, AA. Editing: FC, JS, MRP, CGC, ACJ, MK. Supervision: MV, AA.

## Acknowledgement

This project was financed by DFG VI Grants (908/5-1 and 908/7-1;505660261; 520285844; SPP LOOPS); an NWO VIDI Grant; the Dutch Brain Interface Initiative (DBI2); and by the European Union (ERC Consolidator Grant, NONLIN, 101230764 to MV, and ERC Staring Grant OSCI-PRED, 101075930 to AA). A.C-J. is supported by a Swedish Research Council project grant (VR; 2025-03245), a Research Council of Finland project grant (RCF; 375200), an ANID/FONDECYT Regular (1240899) and ANID/FONDECYT Regular (1251273) research grants. Note that views and opinions expressed are however those of the author(s) only and do not necessarily reflect those of the European Union or the European Research Council. Neither the European Union nor the granting authority can be held responsible for them.

---

**Figure S1:**
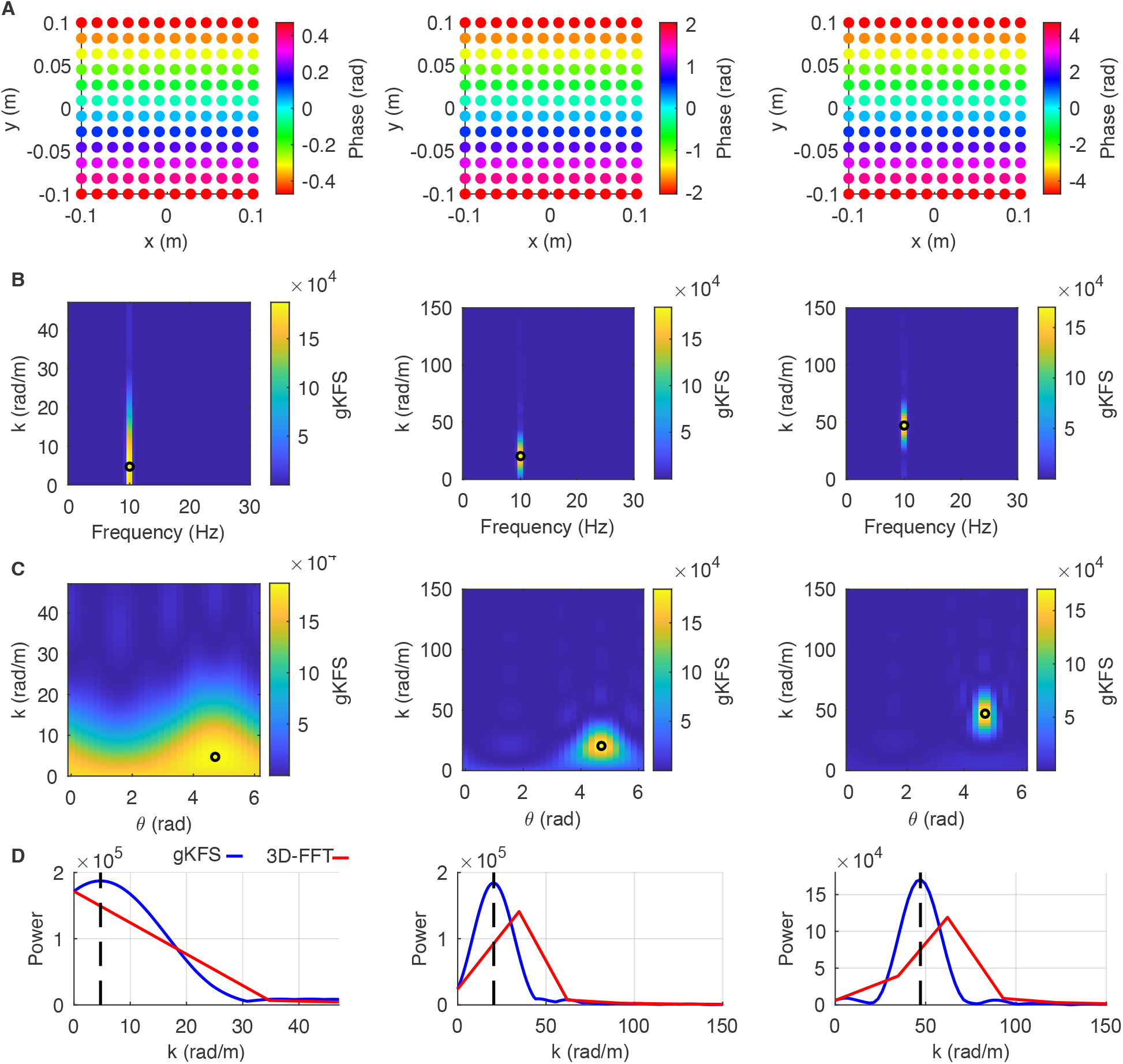
Illustration of the generalized *k*–*f* spectrum. (A) Example of traveling waves at different spatial frequencies. (B) Generalized *k*–*f* spectrum (gKFS) as a function of spatial frequency and temporal frequency (taking the maximum value across wave directions per trial before averaging). The black circle indicates the ground truth wave parameters. (C) The gKFS as a function of wave direction and spatial frequency. (D) Comparison of gKFS (blue) vs. a 3-D Discrete Fourier Transform (red). Note the ability to detect a spectral peak at low frequencies using the gKFS.

**Figure S2:**
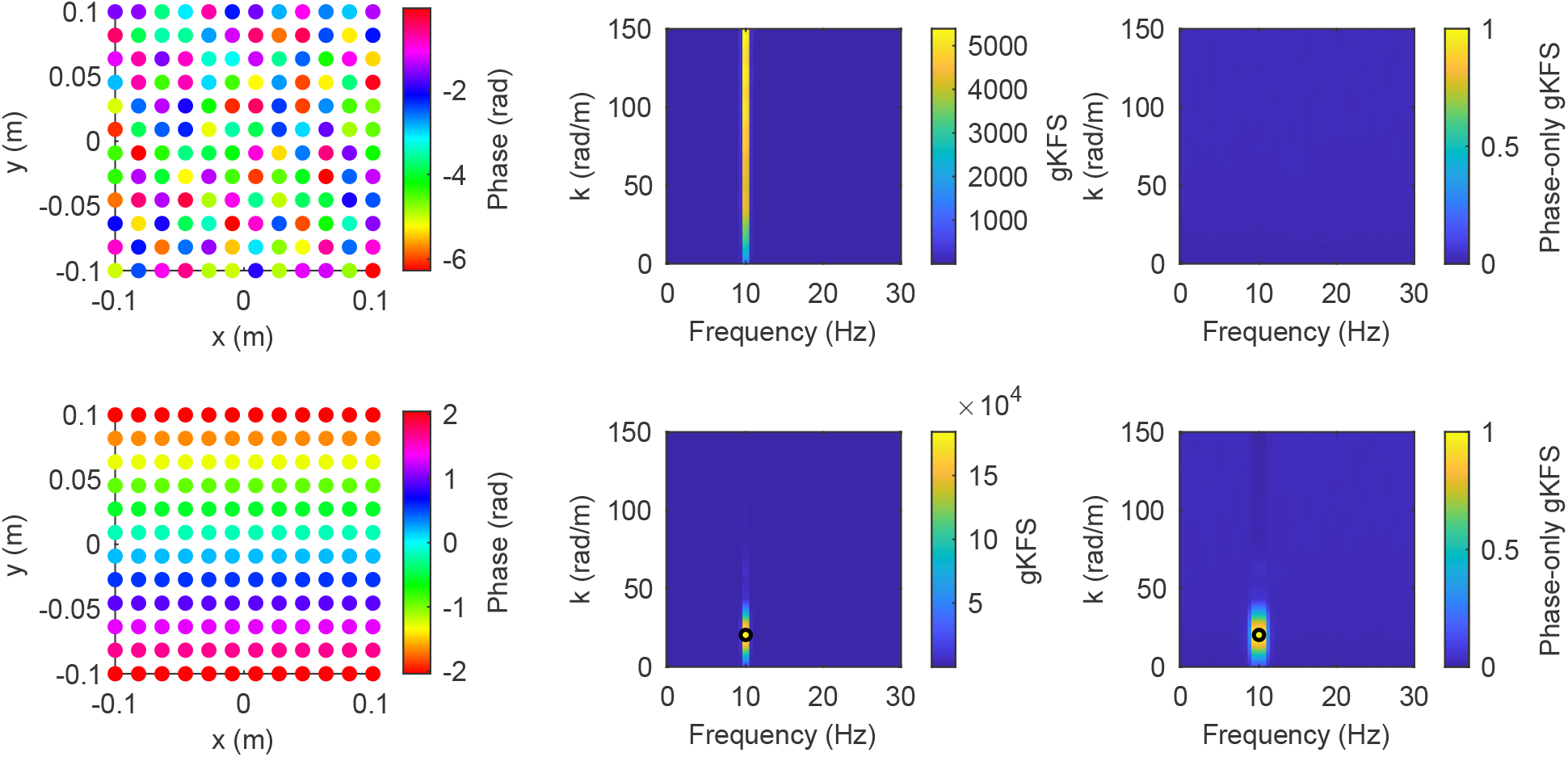
Comparison of generalized *k*–*f* spectrum (gKFS) and the phase-only, amplitude-normalized gKFS (PO-gKFS). Top row: A simulation in which the signal at each electrode is a 10 Hz sinusoid with a random phase, with added white noise. Due to the surplus in power at 10 Hz, the gKFS shows a broad peak at all spatial frequencies. The PO-gKFS does not show any peak due to the lack of phase organization. Bottom row: A simulation in which there is a traveling wave. In this case the gKFS and PO-gKFS show a very similar profile.

**Figure S3:**
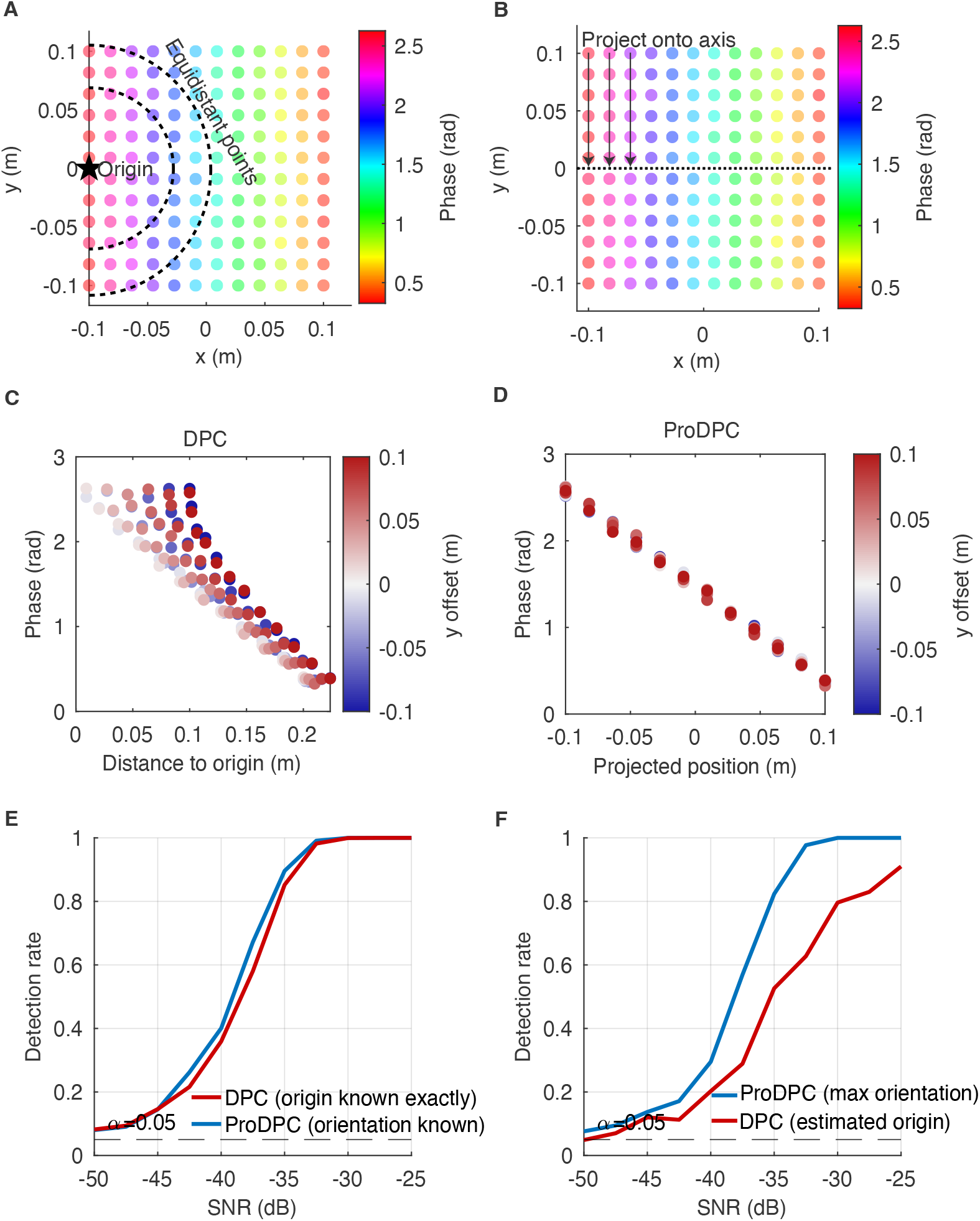
ProDPC vs. DPC for a concrete example: a 12 *×* 12 electrode grid with a planar wave propagating left to right, and a TW origin placed at the left edge of the array. (A) DPC’s algorithm illustrated: the origin (star) and a few iso-distance arcs radiating from it. The arcs only line up with the direction of wave propagation near *y* = 0. (B) Phase map for a single example trial (SNR = 0 dB), with ProDPC’s algorithm illustrated: the propagation axis (dashed line) and a few projection arrows perpendicular to it. Each sensor’s position is converted into a one-dimensional variable by projecting it onto a wave axis; ProDPC then scans through all the wave axes. (C) Distance-to-origin vs. phase for the same trial (extracted from (A)). Points are colored by each electrode’s orthogonal offset *y* from the propagation axis: electrodes with equal phase are assigned different distances purely because of their *y*-position. This results in a reduction of the distance-phase correlation. (D) Projected position vs. phase for the same trial (extracted from (B)). Because projection removes the orthogonal coordinate entirely, there is now a correlation between distance and phase close to one. (E) Single-trial detection rate vs. SNR. In this case, we assume that the true origin is known for DPC and the true wave orientation is known for ProDPC (i.e. we evaluate a best-case-scenario for both methods). Planar waves were generated with varying signal-to-noise ratios. (F) Same as (E), but now a more realistic scenario in which wave direction is unknown (detection statistic taken as the maximum across all candidate propagation angles, with a matched max-based null threshold), and the wave origin is unknown, instead estimated per trial from the point of maximum phase-gradient divergence (Muller et al., 2016). ProDPC shows enhanced detection sensitivity for planar waves compared to DPC.

**Figure S4:**
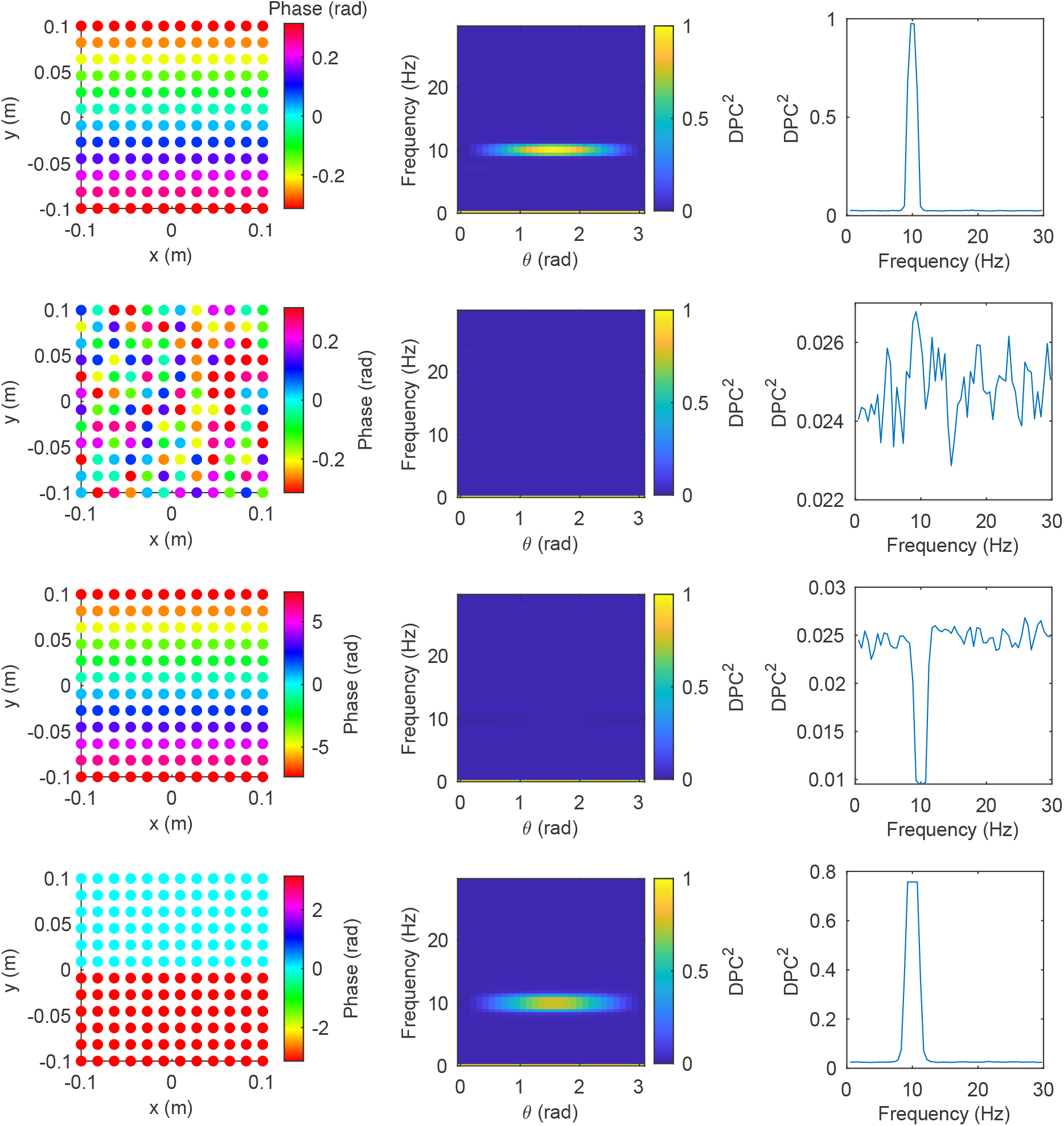
Illustration of properties of the projected distance-phase correlation (ProDPC). First row: Illustration or ProDPC values for a planar wave with a low spatial frequency. Second row: ProDPC for a array with randomly organized phases that has a small range of phases. In this case, ProDPC correctly yields low values. Third row: ProDPC for a planar wave with a high spatial frequency. In this case, ProDPC does not detect the planar wave structure, which is due to the fact it is computed by linearizing phase. Fourth row: ProDPC values for a standing wave. This figure shows that ProDPC does not distinguish between planar and standing waves.

**Figure S5:**
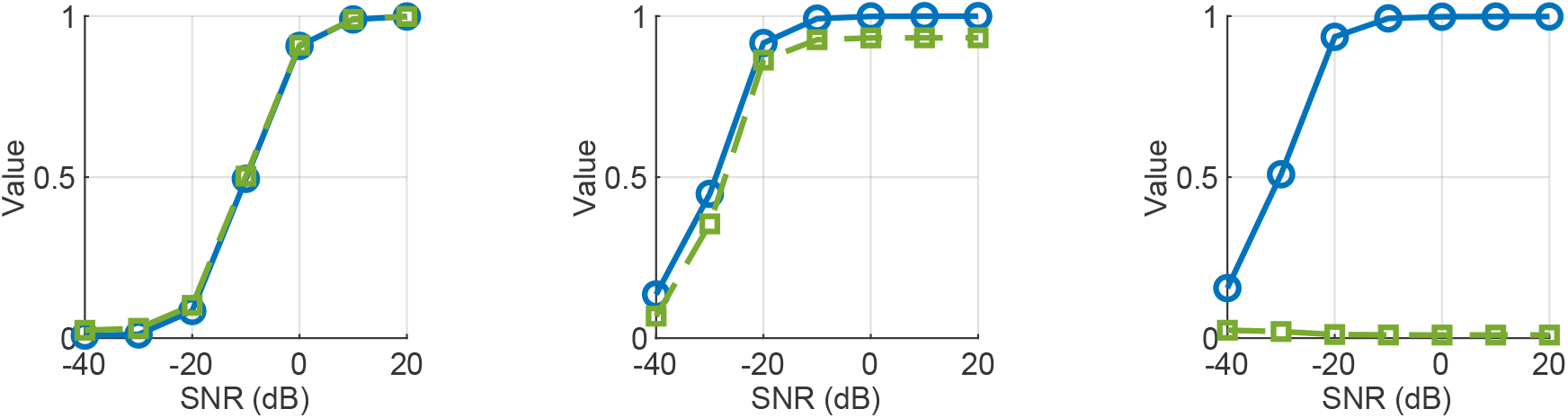
ProDPC (projected distance-phase correlation) (green) and TWINDEX (blue) shown as a function of the signal-to-noise ratio. Left to right: low to high spatial frequencies (0.5, 6.5, 23.5 *π* radians/m). This plot shows that for low spatial frequencies, TWINDEX and ProDPC correlate strongly, reflecting their analytic relationship, however for high spatial frequencies, the relationship is lost, as predicted from the theoretical properties.

**Figure S6:**
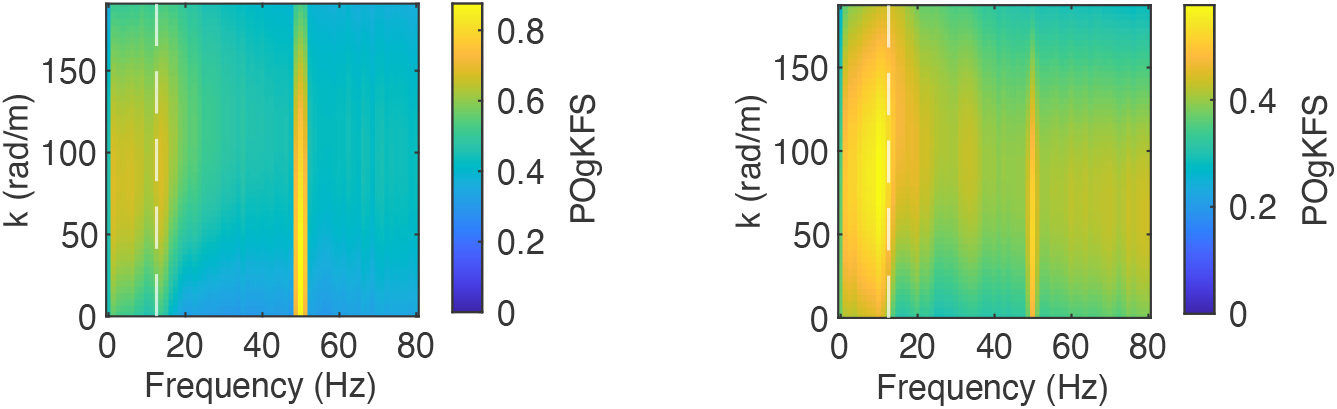
Phase-only gKFS for the marmoset data. This figure shows a relative lack of selectivity across spatial frequencies in the main oscillatory band (12.5 Hz), as compared to the TWINDEX. Furthermore, the figure shows that the PO-gKFS shows a significant peak at the line noise frequency (50 Hz), as opposed to the TWINDEX.

**Figure S7:**
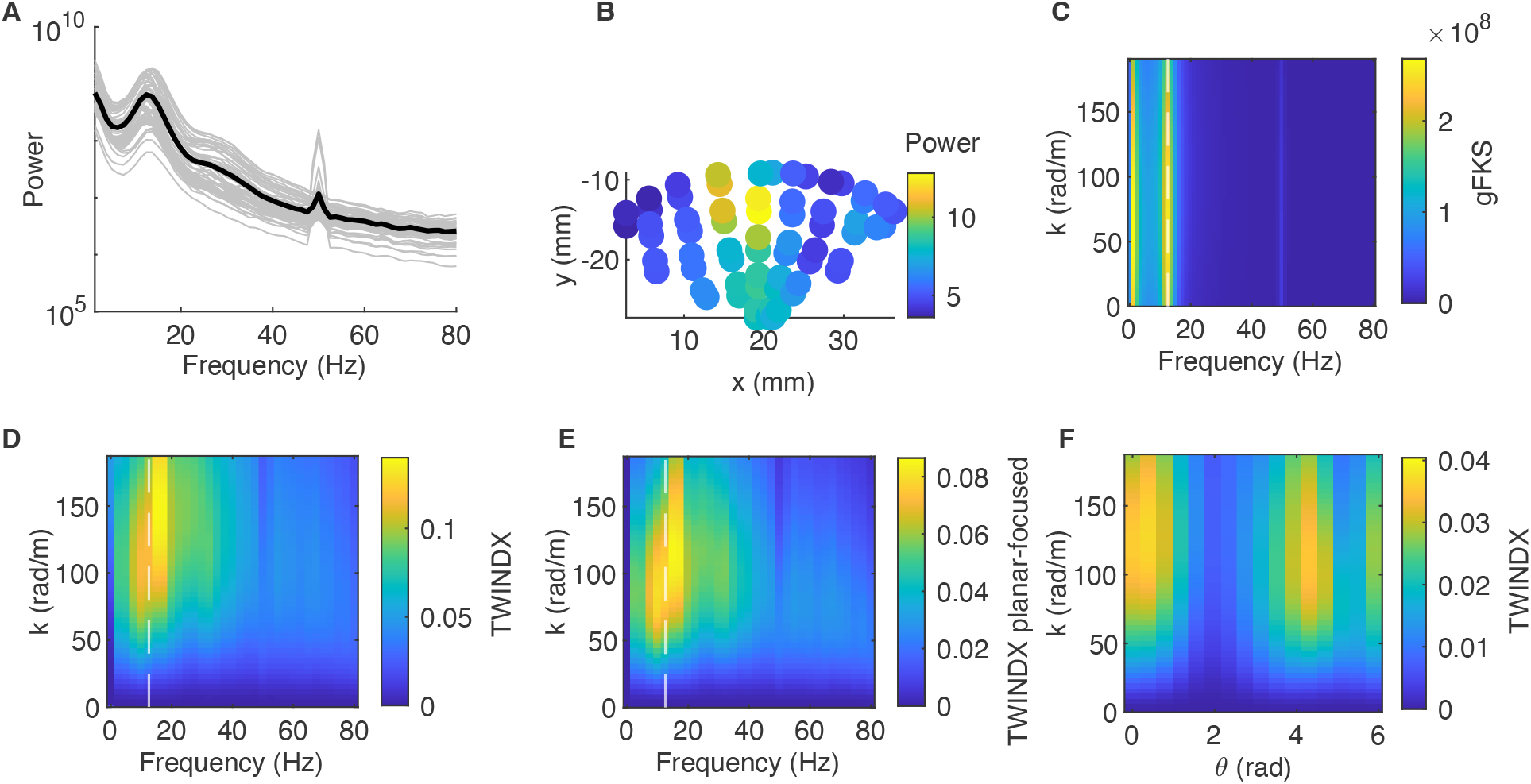
Analysis of ECoG marmoset data during pre-stimulus period, for monkey GO. (A) Power spectra across all channels. Note a genuine oscillatory band around 12.5 Hz, and line noise around 50 Hz. (B) Distribution of power (log-scaled) across channels (left: posterior; right: anterior). (C) Average gKFS spectrum. (D) Average TWINDEX. (E) Average planar-focused TWINDEX. (F) TWINDEX as a function of wave orientation and spatial frequency.

**Figure S8:**
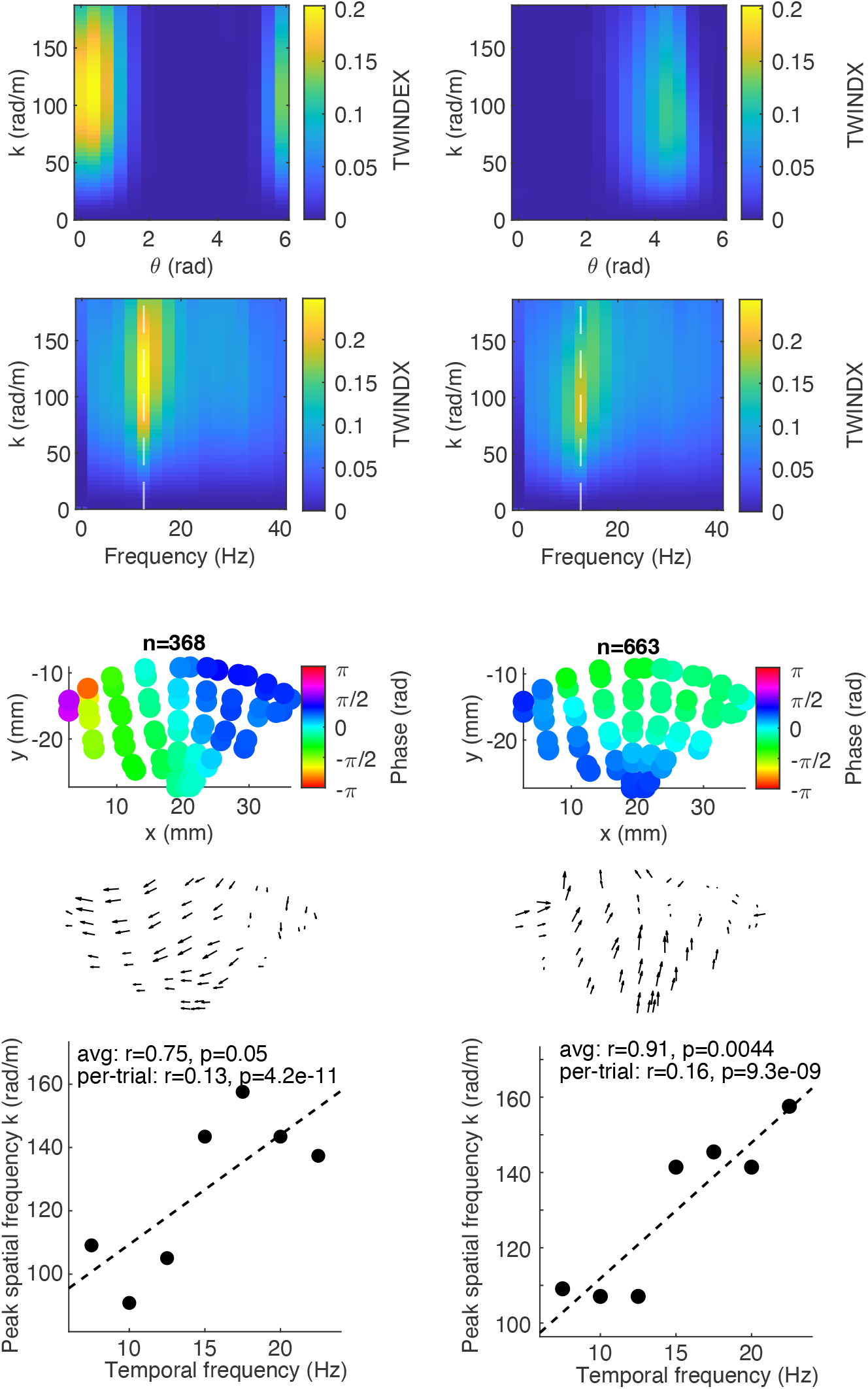
Analysis of separate clusters of trials, for monkey GO. Trials were clustered based on the TWINDEX as a function of spatial frequency and wave orientation, using the HDBSCAN algorithm. Left and right column correspond to two different clusters identified by HDBSCAN. Left column: n=368 trials. Right column: n=663 trials. Top to bottom: TWINDEX as a function of spatial frequency and wave orientation; TWINDEX as a function of spatial and temporal frequency; phase map, obtained by aligning the phases of each trial to a circular mean of zero and then averaging across trials; gradient of the phase map; peak spatial frequency vs. temporal frequency, showing a significant positive correlation both when computing the peak spatial frequency per trial separately, or across trials.

**Figure S9:**
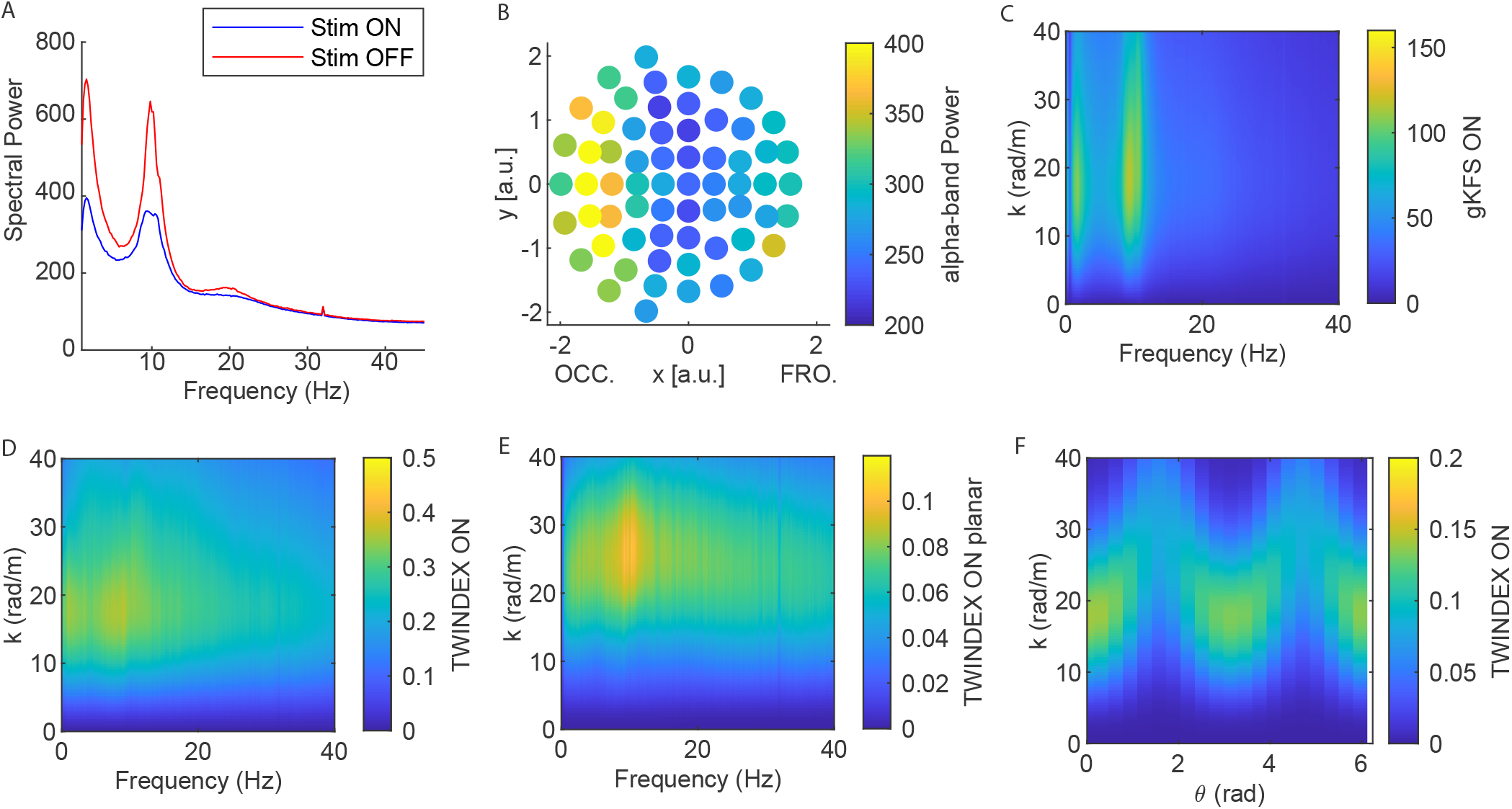
EEG results during visual stimulation. A) Averaged power distribution for all electrodes in the presence (stim ON) and absence (stim OFF) of visual stimulation. B) Topography of the average alpha-band power across participants in the Stim ON condition. OCC. and FRO. denotes the occipital and frontal regions on the x-axis. C,D,E) Generalized *k*–*f* spectrum (gKFS), TWINDEX, and planar-focused TWINDEX applied to scalp EEG data recorded during visual stimulation. F) TWINDEX values as a function of waves’ direction and spatial frequency.

**Figure S10:**
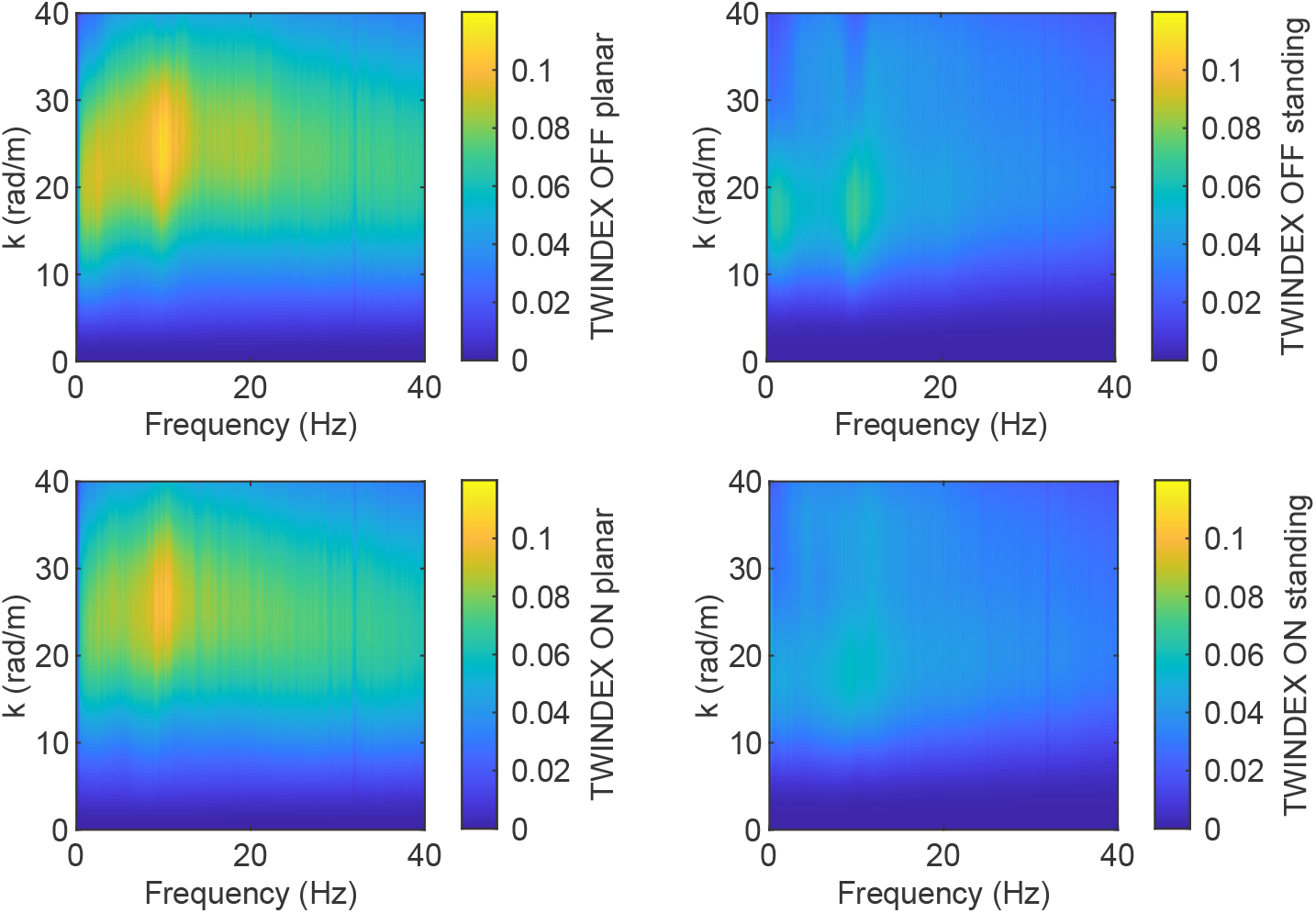
Standing-focused (right panels) and planar-focused (left panels) TWINDEX applied to scalp EEG data recorded in the absence (upper row) and during (lower row) visual stimulation.

